# Heterotypic interfacial tension between oncogenic and wild-type populations forms the mechanical basis of tissue-specific oncogenesis in epithelia

**DOI:** 10.1101/2025.03.14.643229

**Authors:** Amrapali Datta, Phanindra Dewan, Aswin Anto, Tanya Chhabra, Tanishq Tejaswi, Sindhu Muthukrishnan, Akshar Rao, Sumantra Sarkar, Medhavi Vishwakarma

## Abstract

Why does the same oncogenic mutation drive tumor formation in some tissues but not in others? While cancer driver mutations are well-documented, their tissue-specific effects remain largely attributed to genetic factors, leaving the biophysical aspects underexplored. Here, we demonstrate that mechanical interactions, specifically interfacial tension between newly transformed and wildtype epithelial cells are critical in determining survival and growth of HRas^V12^ oncogenic mutants in human mammary and bronchial epithelia, leading to contrasting outcomes in the two tissues. In mammary epithelium, isolated oncogenic cells are extruded-a typical mechanism of defense against cancer in epithelia-while oncogenic groups become spatially confined in a kinetically arrested, jammed state, marked by an actomyosin belt at the interface. In contrast, bronchial epithelium permits persistent spreading of the same oncogenic cells, which form long protrusions regardless of colony size. Furthermore, oncogenic clusters in these two tissues exhibit distinct biophysical properties, including variations in cell shapes, intracellular pressure, cell-cell tension, and cellular motility. Using a cell shape-tension coupled bi-disperse vertex model, we reveal that differences in interfacial tension at mutant–wild-type boundaries dictate whether oncogenic cells are eliminated, restrained, or expanded and that modulating the heterotypic interfacial tension alters mutant cell fate within the epithelium. Together, our findings uncover a mechanical basis for tissue-specific oncogenesis by highlighting how differential cellular mechanics at the oncogenic–host cell interface regulate tumor initiation and progression.

## Introduction

Cancer has traditionally been understood to be a consequence of deregulated cell proliferation driven by genetic alterations that enable clonal expansion and tumor formation ^[1]^. Interestingly, genomic analysis reveals that many tumor suppressor genes and oncogenes exhibit tissue specificity, meaning they are altered in some cancers but not others ^[2-4]^. Current research suggests that a combination of intrinsic biological factors ^[4]^, such as differential epigenetic changes ^[3,5]^ and aneuploidy patterns ^[6-8]^, as well as extrinsic factors like the tumour microenvironment, including the presence of tissue-specific immune cells ^[9-10]^ influences tissue-specific cancer outcomes. However, a relatively underexplored yet crucial aspect of tissue-specific oncogenesis is the mechanical interaction between normal and cancerous cells. To this end, recent studies have highlighted a tumor suppressive behavior in epithelial tissues known as epithelial defense against cancer (EDAC) ^[11]^, where transformed cells are expelled from the tissue by surrounding normal cells ^[12-14]^. However, EDAC can fail ^[14]^, allowing mutant cells to evade extrusion when the microenvironment becomes tumour-permissive, such as in conditions of increased stiffness ^[15]^ or inflammation due to a high-fat diet ^[16]^. The success or failure of EDAC is closely linked to changes in cellular contractility ^[12-14]^ and cell-cell adhesions ^[17]^, suggesting that the underlying biophysical properties of the tissue play a decisive role in determining the outcome of cell competition. Since these properties can vary across different epithelial tissues, tissue-specific differences in cancer susceptibility may be explained by the varying ability of epithelia to manage newly transformed cells. However, despite these advances, the mechanical basis of tissue-specific oncogenesis remains missing. Specifically, how differences in cellular mechanics at the interface between oncogenic and wild-type populations influence tumor initiation and progression is poorly understood. Addressing this gap could not only enhance our understanding of cancer biology but also open new avenues for tissue-specific therapeutic interventions by identifying what makes certain tissues more permissive to tumor formation.

To this end, we investigated and compared the bio-physical factors influencing early tumorigenesis in human mammary and bronchial epithelial monolayers by sporadically transfecting a subset of healthy cells into oncogenic cells expressing a constitutively active HRas^V12^ protein. We then tracked the fate of these transfected cells and their interactions with the wild-type counterparts in real time via live imaging. Strikingly, HRas^V12^-mutant cells exhibited contrasting behaviors in the two tissues. In mammary epithelium, isolated transformed cells were frequently extruded. However, when present in groups, extrusion of oncogenic cells was not seen, but instead, they remained spatially confined in a jammed state. In contrast, in bronchial epithelium, single as well as groups of HRas^V12^ expressing cells persisted and exhibited unrestricted expansion over time, regardless of cluster size. To explain these findings, we employed a bi-disperse vertex model of epithelia ^[18-21]^ and demonstrated that differences in interfacial tension between wild-type and transformed populations play a crucial role in determining whether mutant cells are eliminated, restrained, or are allowed to persist and expand. Taken together, these findings provide novel insights into the differential outcomes of competitive cellular interactions in the precancer stages and uncover a mechanical basis for tissue-specific oncogenesis.

## Results

### 1. Singlets and groups of HRas^V12^ oncogenic mutants show tissue-specific fates in epithelia

To explore the differences in the premalignant stages across epithelial tissues, we sporadically transfected monolayers of the two non-transformed human epithelial cell lines, MCF10A (mammary) and BEAS2B (bronchial), with a plasmid construct that constitutively expresses HRas^V12^ along with the GFP reporter [Figure 1a]. We then monitored the cellular dynamics over 90 hours. Consistent with previous studies reporting extrusion of single HRas^V12^ mutants in MDCK monolayers ^[11-15, 22]^, we observed that isolated HRas^V12^ transfected cells (singlets) surrounded by wild-type neighbours were extruded from the mammary epithelium [Figure. 1b (upper panel), Supplementary video 1]. This process correlated with a reduction in shape indices of oncogenic mutants as the density of wild-type cells around them increased [Figure. 1c (upper panel)]. Interestingly, this well-known mechanism of epithelial defense against cancer was impaired in BEAS2B cells, where HRas^V12^ mutants evaded extrusion [Figure. 1b (lower panel), d], and wild-type cell compaction did not restrict their growth [Figure. 1c (lower panel), Supplementary video 1]. Instead, the mutants in the bronchial epithelium continued to proliferate and exhibited increased elongation with long protrusions [Figure. 1c (lower panel), e, Supplementary video 1]. Given that cancers frequently arise from fields of oncogenic mutants rather than isolated mutated cells ^[23-25]^, we also investigated whether these differential dynamics of mutant singlets would also extend to groups of oncogenic mutants, potentially enabling more aggressive HRas^V12^ growth in bronchial tissue compared to mammary tissue. By adjusting cell seeding conditions, we generated larger groups of HRas^V12^ transfected cells in both MCF10A and BEAS2B monolayers and then tracked the dynamics of these groups over time. In the mammary epithelium, wild-type cells were unable to extrude groups of HRas^V12^-transfected cells but instead confined them into compact, circular clusters with smooth, rounded interfaces [Figure 1f, (upper panel), g (upper panel), Supplementary Video 2]. As a result, oncogenic clusters gradually demixed from the surrounding wild-type MCF10A population and remained spatially constrained [Figure 1h]. In contrast, HRas^V12^ clusters in the bronchial epithelium gradually spread outward [Figure 1f, (lower panel), g (lower panel), Supplementary video 2], and formed protrusive lamellipodia and developed an irregular interface with surrounding wild-type cells [Figure. 1g (lower panel)]. Together, these striking differences in the dynamics of HRas^V12^ oncogenic cells demonstrate that activation of the same oncogene can lead to distinct outcomes across tissues in terms of growth, spreading, and extrusion. This prompted us to further investigate the unique bio-physical mechanisms driving these variations.

**Figure 1:**
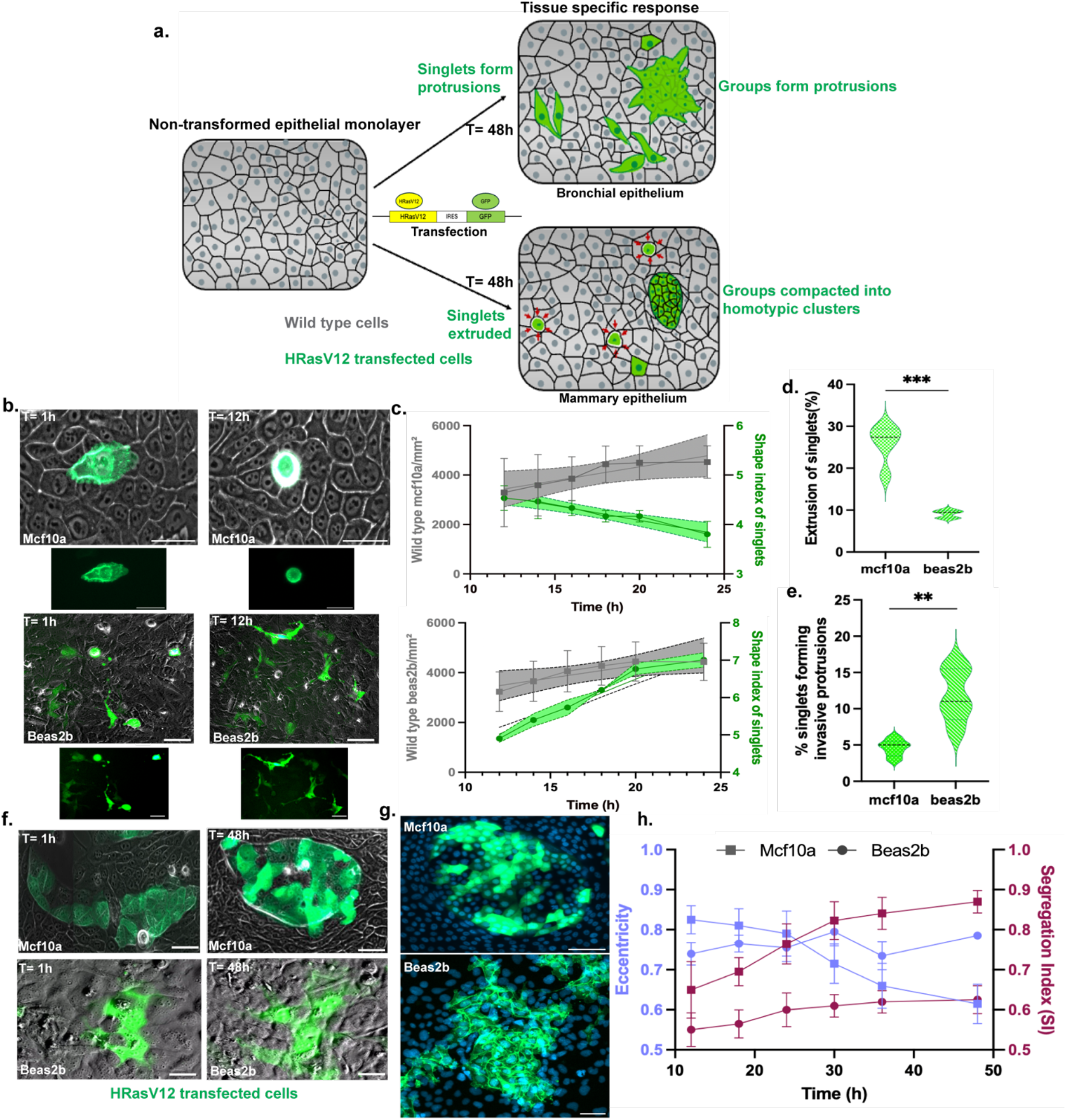
Tissue-specific outcomes of HRas^V12^ oncogenic mutants in epithelial monolayers. (a) Schematic representation of the experimental setup and outcome (b) Representative images showing extrusion of single oncogenic cells (singlets) in mammary epithelium (*upper panel*), but protrusive behavior of the same oncogenic singlets in bronchial epithelium (*bottom panel*) (c) Quantification of shape indices of HRas^V12^ singlets showing a reduction in shape indices, with increasing wild-type cell density in mammary epithelium (*upper panel*), but the opposite trend in bronchial epithelium (*bottom panel*), where HRas^V12^ cells continues to spread. Shaded regions indicate the results simple linear regression analysis with 95% confidence intervals (d) Extrusion rates of HRas^V12^ singlets are significantly higher in mammary epithelium, while (e) Percentage of HRas^V12^ cells forming protrusions is markedly higher in bronchial epithelium. Representative data are plotted from one of three independent experiments with the median shown as a bold dashed line and the first and third quartiles are shown as thin dashed lines. Statistical significance was calculated using Unpaired t-test with Welch’s correction. (f) Behaviour of HRas^V12^ oncogenic clusters in the two tissues-In mammary epithelium (*upper panel*), clusters become spatially confined with a smooth, circular interface with the wild-type population, while in bronchial epithelium (*bottom panel*), HRas^V12^ clusters expand, forming long protrusions over time (g) Immunofluorescence images of HRas^V12^ clusters in mammary and bronchial epithelia, highlighting differences in spreading patterns in the two epithelia (h) Quantification of cluster segregation: mammary epithelium exhibits a higher segregation index and reduced eccentricity compared to bronchial epithelium, indicating greater spatial confinement of oncogenic clusters. The segregation index (SI), defined as the average ratio of homotypic and all cell neighbors, quantifies the degree of demixing and is averaged over all cells inside an oncogenic cluster. Data are mean±sem and plotted from different clusters from one representative experiment. (*Scale bars= 50 μm*)

### 2. Biophysical analysis reveals demixing via jamming of oncogenic clusters in MCF10A but unjamming and protrusive growth in BEAS2B

To understand the mechanisms driving the differential fates of oncogenic fields in the two epithelial tissues, we mapped the bio-physical signatures of wild-type-mutant interactions by subdividing the images into four regions of interests for both monolayers-ROI1: wild type cells distant from oncogenic cells, ROI2: interfacial wild type cells in direct contact with oncogenic cells, ROI3: oncogenic cells at the cluster boundaries in direct contact with wild type cells, and ROI4: oncogenic cells in contact with other oncogenic cells [Figure. 2a]. We then compared the F-actin levels and shapes of both mutant and wild-type populations across these ROIs. Additionally, we employed Bayesian force inference, as previously established ^[26]^, to infer the differential tissue stresses associated with cluster formation from the observed cell geometries. Interestingly, the differences across these parameters were apparent not only between wild-type and oncogenic counterparts but also based on the spatial localization of the cells, with cells at the oncogenic-wild type interface exhibiting distinct behaviors, regardless of their origin.

**Figure 2:**
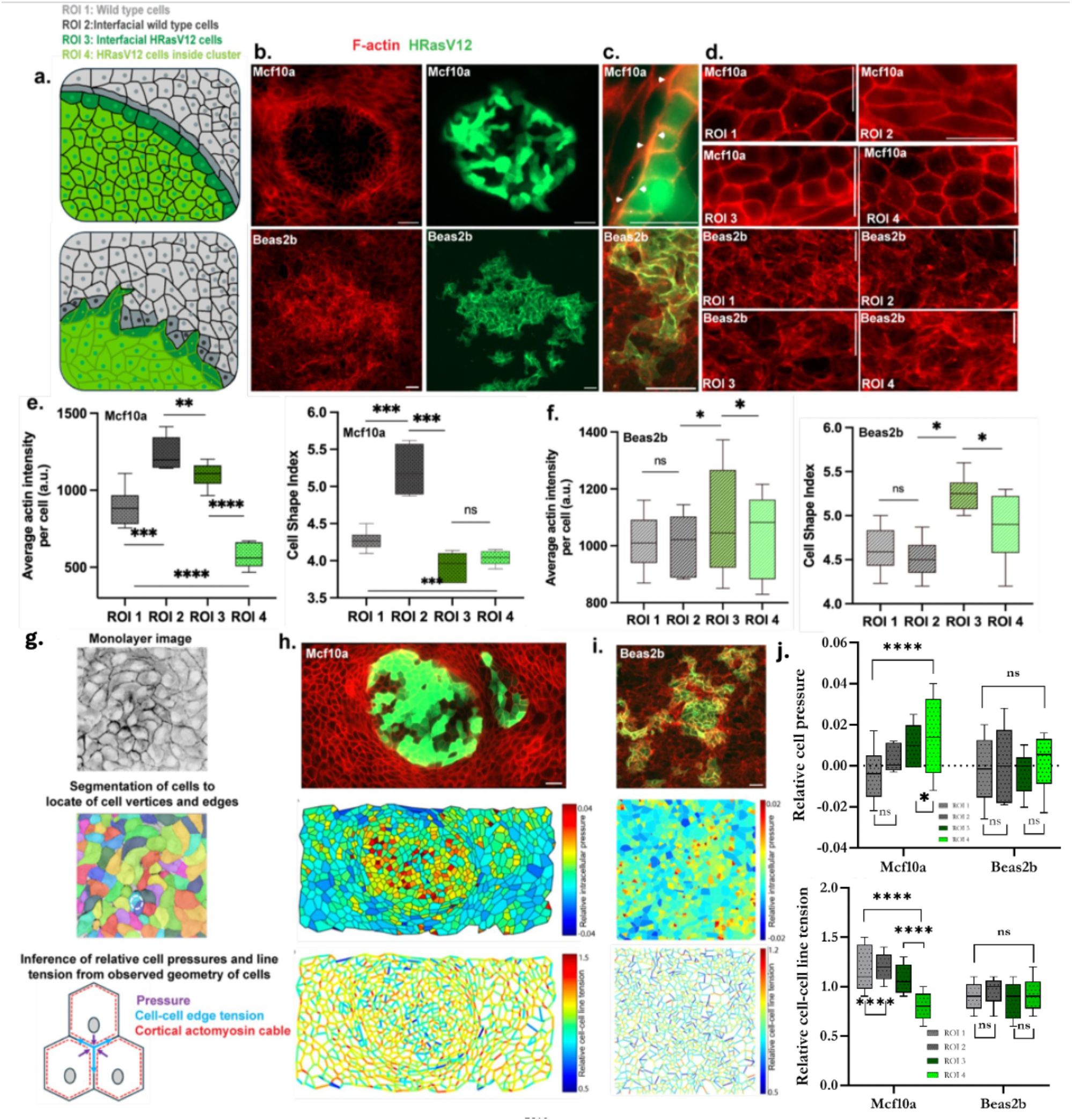
Distinct mechanical signatures of HRas^V12^ clusters in mammary and bronchial epithelia. (a) Representative images showing regions of interest (ROIs) selected for analysis in mammary (*upper panel*) and bronchial epithelia (*lower panel*) (b) Immunofluorescence images of HRas^V12^ clusters stained for F-actin in mammary (*upper panel*) and bronchial epithelia (*lower panel*) (c) Oncogenic cluster–wild-type interface showing distinct actin belt in mammary marked by white arrows (*upper panel)* and absence of it in bronchial epithelia (*lower panel)* (d) ROI-based F-actin stained regions of wild-type and oncogenic cells in mammary (upper panel) and bronchial epithelia (lower panel) revealing distinct shape differences in different regions-in mammary epithelia, wild type cells at the interface show elongated shapes (ROI2), and oncogenic cells inside the cluster show jammed shapes (ROI4), in comparison to wild type cells at a random location (ROI1), while in bronchial epithelia, oncogenic cells (ROI 3 and 4) show more elongated shapes compared to wild-type beas2b cells (e), (f) Quantifications of F-actin intensities and shape indices, in mammary epithelia (e), and bronchial epithelia (f), across the four ROIs confirming jammed oncogenic clusters, with unjamming at the interface, in mammary epithelia and unjammed clusters in bronchial epithelia (g) Schematic illustrating the Bayesian Force Inference pipeline used to estimate relative intracellular pressure and cell–cell edge tension (h) Heatmaps depicting relative cell pressures and cell–cell edge tensions in mammary epithelium, revealing localized mechanical heterogeneity (i) Heatmaps of relative cell pressures and edge tensions in bronchial epithelium, showing a distinct mechanical landscape compared to mammary epithelium (j) Quantification of relative intracellular pressures and cell–cell edge tensions across the four ROIs in mammary (upper panel) and bronchial epithelia (lower panel) highlighting tissue-specific differences. All data represented as mean±sem in box and whisker plots and are plotted from one out of three independent experiments with whiskers extending from the minimum to the maximum values, showing the full range of the dataset including outliers. Line represents the median. Statistical significance was calculated using Unpaired t-test with Welch’s correction. *Scale bars= 20 μm*

In mammary epithelium, oncogenic clusters were surrounded by a contractile actomyosin belt [Figure. 2b (upper panel), c (upper panel)], shown by a prominent deposition of F-actin and phosphomyosin at heterotypic (oncogenic cluster-wild type population) interfaces [Supplementary Figure 4] coupled with drastically low F-actin levels inside the clusters [Figure. 2b (upper panel), e (left panel)], suggesting that the de-mixing of oncogenic cells in mammary tissue is likely driven by changes in cytoskeletal mechanics. Further, oncogenic clusters in mammary epithelium (ROI4) also exhibited low cell shape indices [Figure. 2d (upper panel), e (right panel)], high internal pressure and low line tension [Figure. 2h, j]-characteristics indicative of jamming ^[27]^. In epithelial tissues, the jamming–unjamming transition is closely linked to changes in cell shape ^[27]^ and is often quantified using the cell shape index (the ratio of cell perimeter to the square root of area), where lower shape indices correspond to jammed, solid-like states, and higher shape indices reflect unjammed, fluid-like tissue behavior. Notably, both oncogenic cells and wildtype MCF10A cells at the cluster interface (ROI 2 and ROI3) showed F-actin enrichment, while the interfacial wild shape cells show elongated shapes [Figure. 2d (upper panel), e], indicating that interfacial cellular mechanics play a key role in the de-mixing process as the wild-type cells encircle the oncogenic mutants. This interfacial region enriched with F-actin also contained phosphomyosin [Supplementary Figure. 1a], indicating that a contractile actomyosin belt drives the compaction of oncogenic clusters in mammary epithelia. In contrast, bronchial epithelium exhibited no actin belt formation at the wild type-oncogenic cell interface, and no traits of jamming were observed in the oncogenic clusters [Figure. 2b (lower panel), c (lower panel)]. Instead, oncogenic cells continued to proliferate, displaying elongation measured via higher shape indices and high F-actin expression [Figure. 2d (lower panel), f]-indicative of unjamming behaviour ^[27,28]^. The relative internal cell pressure and cell-cell line tension inferred with Bayesian force inference showed slight or no modulation in the oncogenic clusters, given the general elongated shapes of wild-type cells in BEAS2B cells [Figure. 2i, j]. Together, these findings highlight distinct interfacial mechanics between oncogenic cells and their wild-type neighbors in both MCF10A and BEAS2B monolayers, resulting in differing outcomes in mutant colony growth.

### 3. PIV analysis reveals kinetic arrest of oncogenic clusters in MCF10A coupled with tangential motion of interfacial cells

To further probe the cellular dynamics that lead to the contrasting fates of the oncogenic mutant colonies in the two tissues, we analyzed cellular velocities during the de-mixing of oncogenic cells, using Particle Image Velocimetry (PIV) on consecutive image pairs from mammary and bronchial monolayers. Interestingly, velocity maps in samples imaged for mutant cluster – wild type dynamics in MCF10A [Supplementary video 3] revealed an unusual movement pattern at confluency: wild type MCF10A cells exhibited tangential motion around oncogenic clusters [Figure. 3c], which coupled with a drastic reduction in the velocity of oncogenic cells [Figure. 3a (lower panel), d], likely caused their compaction and jamming into clusters. Notably, this tangential movement was unique to wild-type MCF10A cells surrounding oncogenic clusters and was absent in other regions across the monolayer [Figure. 3c, Supplementary Figure 5] as well as under control conditions-i.e., in the absence of oncogenic mutants [Supplementary video 4], in both of which a typical velocity drop was observed in MCF10A cells due to density-induced jamming ^[27-30]^ [Figure. 3d (upper panel), Supplementary Figure 2]. These results aligned with the observed differences in cell shape indices, where oncogenic clusters underwent jamming, while wild-type cells at the interface exhibited unjamming behavior ^[29,30]^. In contrast, in the bronchial epithelium, mutant clusters remained unjammed, even as surrounding wild-type cells became more crowded [Figure. 3b, d (lower panel), Supplementary video 5]. Additionally, no tangential cellular movement was detected around the oncogenic clusters in the bronchial epithelium. These velocity maps showed negligible deviations from wild type (control) BEAS2B velocity maps [Supplementary video 6]. Together, these findings suggest that mutant clusters carrying the same oncogene can exhibit drastically different behaviors depending on the epithelial tissue type. Specifically, differences in the dynamics of the interfacial cells drive the demixing and jamming of HRas^V12^ oncogenic cells in mammary epithelium while promoting protrusive, unjammed growth in bronchial epithelium.

**Figure 3:**
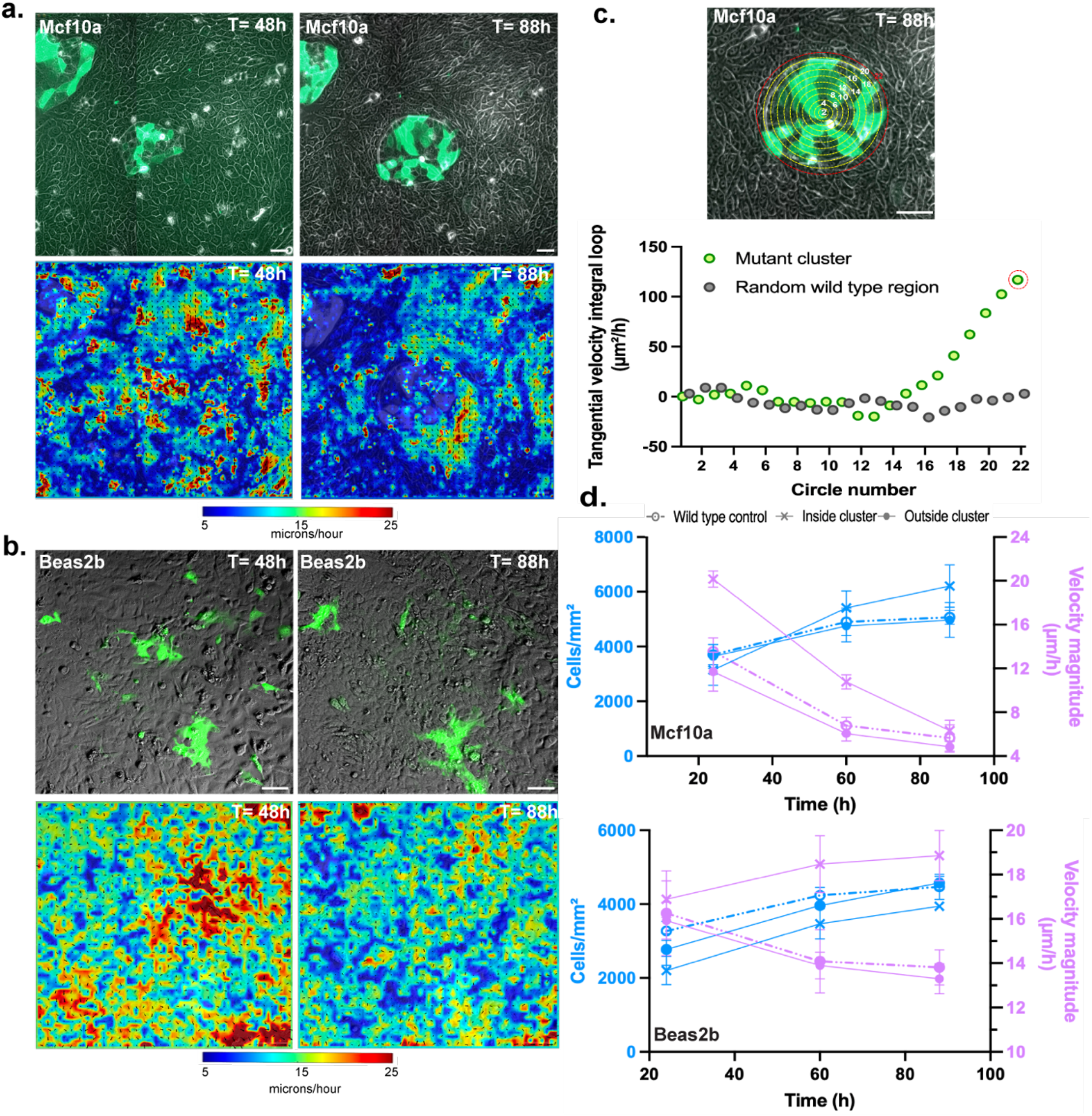
Particle Image Velocimetry (PIV) reveals distinct cellular movement patterns in HRas^V12^ clusters in the two monolayers. (a), (b) Representative snapshots of HRas^V12^ clusters, and the corresponding PIV maps, in mammary (a) and bronchial epithelium (b), as the density of wild type cells increase showing wild type cells along the cluster-wild-type interface exhibit a higher velocity attributed to tangential motion in mammary epithelium (a), but undergo a more kinetically arrested state in bronchial epithelium (b) (c) Oncogenic cluster in MCF10A monolayer with concentric circles drawn from the centre of the cluster, radially outward till its boundary (upper panel) with the red (highlighted) circle representing the one along which tangential velocity was the highest and plot showing tangential motion (expressed as a function of the circles), revealing highest tangential motion (data point highlighted with red) along the interface of oncogenic cluster with the wild-type cells in mammary epithelia (lower panel). No significant tangential motion in random wild-type region (c) Representative snapshots of HRas^V12^ clusters in bronchial epithelium at the same two time points and the corresponding PIV velocity maps (d) Quantification of cellular velocities in local regions as cell densities reach confluency showing oncogenic cells clustered in mammary epithelium (*top panel*) jam and show a significant reduction in velocities. In bronchial epithelium (*bottom panel*) while wild type cells show reduced movement as cell density increased, transfected cells continued to show higher motion, signifying that jamming of wild type cells has no effect on mutants. All data are represented as mean±sem plotted from different clusters from one representative experiment. *Scale bars= 50 μm*.

### 4. Bi-disperse vertex model reveals interfacial tension between mutant and wild-type population governs tissue-specific response towards oncogenic mutants

To explain the differential dynamics and degrees of demixing of oncogenic clusters observed in the two tissues, we drew inspiration from the differential interfacial tension hypothesis (DITH) ^[31]^. This hypothesis suggests that cell sorting in tissues could occur due to the variations in interfacial tension, as seen in embryonic cell rearrangement, where cells regulate contractility, as well as adhesions to demix from their neighbors ^[31-33]^. To test whether such a mechanism could be applied to the de-mixing of oncogenic clusters from wild-type cells-leading to tissue-specific outcomes, we employed the cell-based bi-disperse Vertex Model (VM). Our model allowed us to simulate an epithelial monolayer and capture the dynamic interactions between wild-type and mutant cell populations. We analyzed the mechanics underlying the segregation dynamics of mutant clusters by examining the interfacial line tension 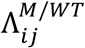 between the oncogenic and wild-type cells [Figure. 4a]. In our model, Λ_*ij*_ = 0, for homotypic interfaces, i.e. wild-type-wild type or mutant-mutant interactions, and Λ_*ij*_ = Λ ≠ 0, for heterotypic interfaces between oncogenic and wild type cells. When Λ > 0, our model ensures that the interface between oncogenic and healthy cells contracts to minimize energy, leading to a reduction in total interfacial length, and when Λ < 0, our model promotes a rough interface by favouring increased interface between healthy and oncogenic cells [Supplementary Video 7]. The final interface shape is determined by the balance between interfacial tension and the intrinsic perimeter and area elasticity of individual cells [Figure. 4b].

**Figure 4:**
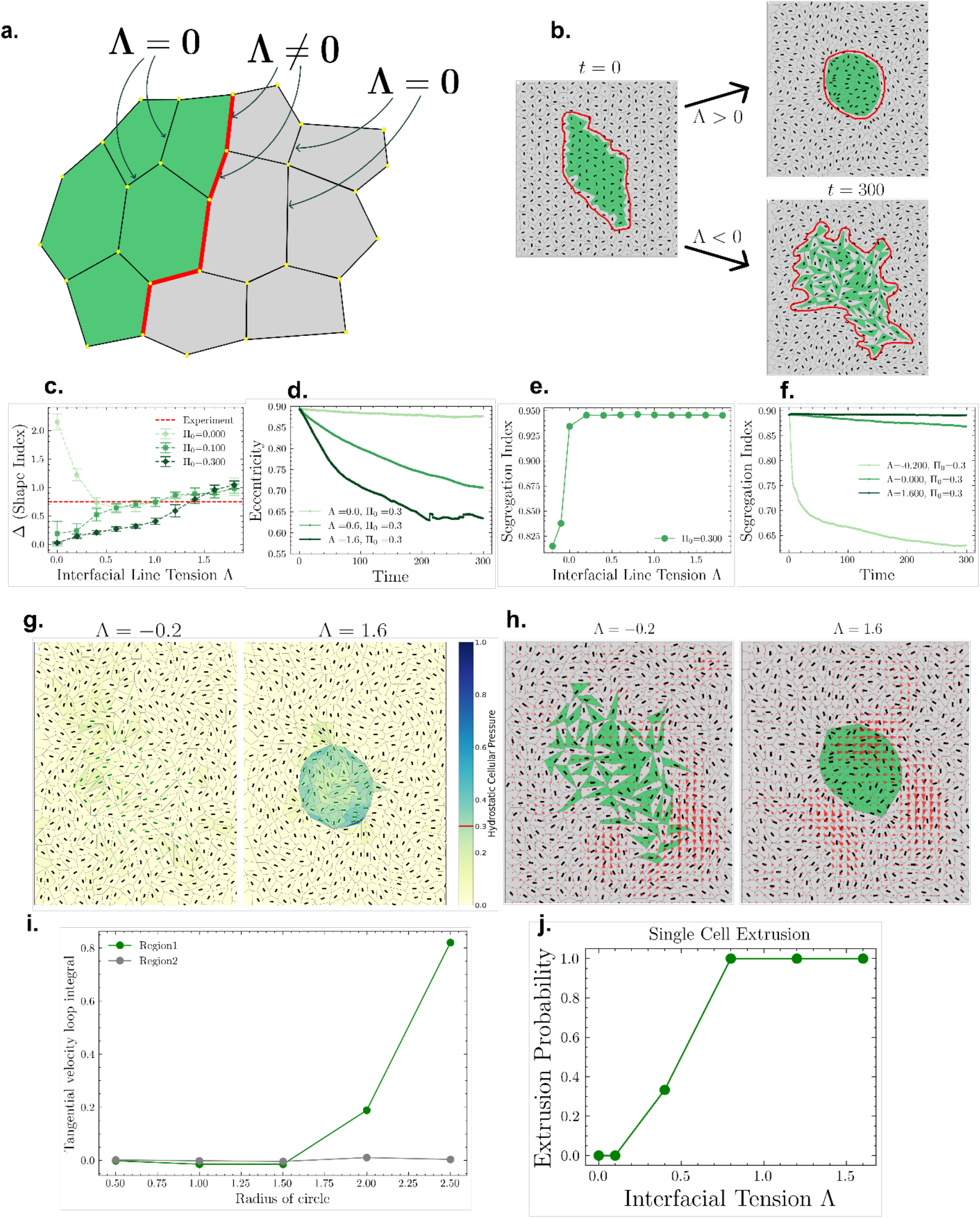
Bi-disperse vertex model reveals interfacial tension-driven segregation of oncogenic clusters. (a) Schematic of the bi-disperse vertex model, where interfacial line tension (red) is applied at the boundary between mutant (green) and wild-type (grey) (b) The value of Λ determines whether the mutant cluster remains compact or spreads into the wild-type population, where the red line gives the outline of the morphology of the mutant cluster interface (c) Difference in shape index between interfacial and bulk cells as a function of Λ (interfacial tension), shown for different stress thresholds. Experimental average (red) overlaid for comparison (d) Eccentricity of mutant clusters (given by the red lines in (b)) over time for different values of Λ (with Π_0_ = 0.3), indicating changes in cluster morphology (e) Segregation index at t = 300 as a function of Λ forΠ_0_ = 0.3, quantifying the extent of cluster segregation (f) Time evolution of the segregation index for different values of Λ (with Π_0_ = 0.3), demonstrating interfacial tension-driven segregation (g) Hydrostatic pressure maps of mutant and wild-type cells at t = 300, with mutant cells labeled by green directors and wild-type cells by black directors (h) Velocity field maps of epithelial monolayers for different values of Λ, revealing flow patterns around oncogenic clusters (i) Tangential velocity loop integral for circles of varying radius centered at the mutant cluster (Region 1) and centered away from the cluster (Region 2), showing tissue-specific velocity variations (j) Extrusion probability of a single mutant cell as a function of Λ, demonstrating how interfacial tension influences mutant cell fate.

To explain the observed differences in interfacial cellular dynamics, we introduced an active contribution to interfacial tension, which was coupled to the shape of cells at the interface, called shape-tension coupling^[34]^. This active interfacial tension is extensile, meaning it elongates cells along the interface depending on their orientation relative to the interface. It originates from the anisotropic distributions of stress-fibres in when cells are under external stress. Hence, we assume that it becomes significant only when the hydrostatic pressure on the interfacial cells exceeds a threshold value Π_0_ [Figure. 4b, g].

Remarkably, our model robustly captured the experimentally observed behaviors. As a first check, we compared the experimentally observed differences in the shape indices between interfacial and non-interfacial wild-type and oncogenic cells in both mammary and bronchial epithelium with our model’s predictions. For Γ = 0.2, Π_0_ = {0, 0.1, 0.3}, Λ = {0 − 1.6}, our model accurately captured the experimental observations [Figure. 4c]. For stress threshold values lower than Π_0_ = 0.3, we find that the differences in shape indices are non-zero even for Λ = 0, which is an unphysical situation. This is unphysical because we expect there to be no difference in the shape indices because in principle, there is only one cell type for Λ = 0. Thus, we have taken Π_0_ = 0.3 for further analyses. To make the comparison between the model and the experiment more robust, we compared other metrics as well.

To quantify the extent of mixing between wild-type and mutant cells, we used the segregation index. Our simulations showed that when Λ < 0 the segregation index rapidly decreases over time, indicating outward protrusion of the mutant cluster. In contrast, for Λ > 0 the segregation index remained close to 1, reflecting a compact, jammed cluster [Figure. 4f]. Additionally, we observed that when Λ > 0, cells within the cluster experience higher internal pressure compared to when Λ < 0 [Figure 4g], consistent with experimental findings [Figure. 2h,i]. Line tensions along edges in the vertex model have also been calculated [Supplementary Figure 6]. Furthermore, flow field analysis from our simulations [Figure 4h,i] revealed that for Λ > 0, cell movement aligns tangentially along the cluster boundary—consistent with the MCF10A behaviour—whereas for Λ < 0 cell motion was uniform throughout the monolayer, resembling the BEAS2B phenotype [Figure. 3a, b]. These comparions further establish that different values of Λ correspond to different cell types. The same oncogene thus might affect different cell lines differently. We also estimated the probability of single mutant cell extrusion as a function of Λ [Figure. 4j], consistent with the findings from our experiments which show that isolated HRas^V12^ transfected cells are frequently extruded in mammary tissue [Figure. 1b]. Further, to check how actin distribution in the tissue can change in response to the interfacial tension in the model, we performed simulations of a vertex model with interfacial tension and mechanochemical feedback ^[35]^. The mechanochemical feedback incorporates a load-dependent myosin-binding mechanism, allowing cells in the vertex model to adjust their fractions of stress fibers and cortical actin individually. The simulation snapshots matched quite well with those from experiment [Appendix figure 5]. Together, our findings demonstrate how interfacial tension-driven mechanics could play a crucial role in determining the competitive dynamics between wild-type and mutant populations, thus providing compelling evidence that cellular and tissue mechanics are key determinants of cell competition.

**Figure 5:**
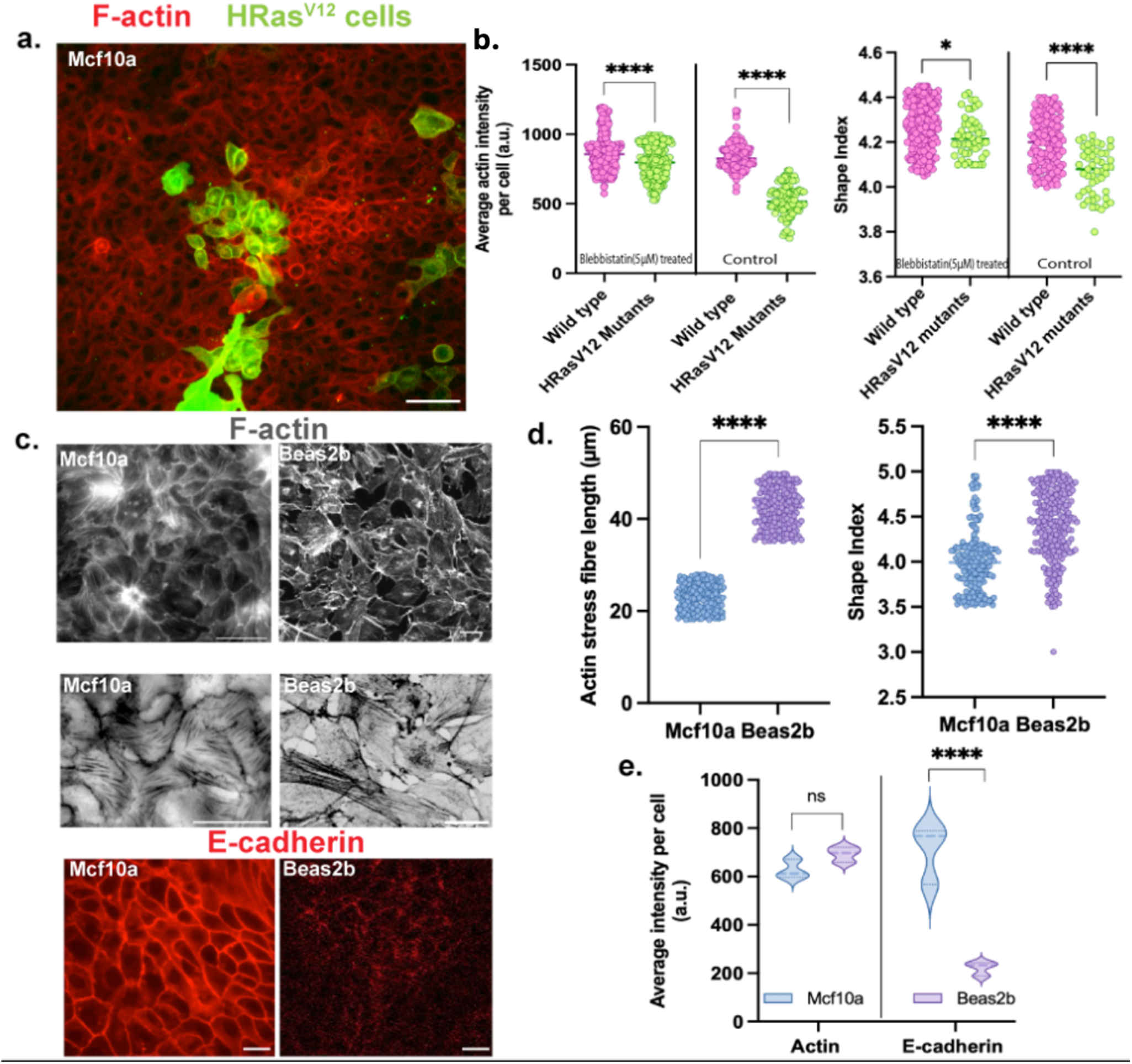
Actomyosin belt disruption prevents mutant cluster compaction in mammary epithelia. (a) Representative actin staining image upon blebbistatin treatment, done after oncogenic induction in mammary epithelial monolayer, showing an absence of an actin belt (b) Quantification of F-actin intensities (left) and shape indices (right), showing difference between oncogenic clusters and wild-type cells in these parameters, is getting small – although not completely vanished because of low concentration of blebbistatin used for the experiment (c) Representative actin and E-cadherin staining images of the wild type monolayers showing shorter stress fibers, higher E-cadherin intensities in MCF10A wild-type monolayer, and the opposite in BEAS2B (d) Plots comparing F-actin stress fiber length (left panel) and shape indices (right panel) in wild-type MCF10A and BEAS2B monolayers. Stress fiber lengths were calculated manually in ImageJ (e) Plots comparing F-actin and E-cadherin levels in the two wild-type tissues. Representative data are plotted from one of three independent experiments with the median shown as a bold line. Statistical significance was calculated using Unpaired t-test with Welch’s correction. *Scale bars= 50 μm*.

### 5. Disrupting actomyosin belt using blebbistatin prevents demixing and compaction of mutant clusters in mammary epithelia

Since our model showed that differences in interfacial tension regulate mutant cluster dynamics, with a higher interfacial tension driving cluster compaction [Supplementary video 7], we reasoned that this tension is mediated by the supracellular actomyosin belt ^[36]^ that drives the compaction and segregation of oncogenic clusters in mammary epithelium. Therefore, we investigated whether disrupting the cytoskeletal organization and impairing actomyosin belt formation would affect the demixing and compaction of oncogenic clusters. To test this, we treated MCF10A monolayers with 5 µM blebbistatin, a myosin II inhibitor, to disrupt belt formation.

As expected, blebbistatin treatment prevented mutant cluster compaction [Figure. 5a], leading to a loss of their rounded morphology and restoring actin levels and shape indices of mutant cells to wild-type levels [Figure. 5b]. This shift resulted in mutant clusters resembling those observed in bronchial epithelia, suggesting that cytoskeletal organization plays a key role in tissue-specific differences in mutant cluster dynamics. This was also consistent with the fact that interfacial tension between cells arises from the interplay between the cortical actomyosin network and cell-cell adhesion ^[37]^.

Additionally, we looked into the inherent differences in cytoskeletal mechanics of the host tissues and observed distinct differences in actin organization between the two tissues. Wild-type bronchial and mammary epithelial cells exhibited comparable F-actin intensities [Figure. 5c (upper panel), e], although the bronchial epithelial cells had longer stress fibers [Figure. 5c (middle panel), d (left panel)]. These cytoskeletal differences were also coupled with significant differences in cell shapes between confluent MCF10A and BEAS2B monolayers with wild-type BEAS2B cells exhibited significantly higher shape indices, indicating elongated, loosely packed morphologies and a lower degree of jamming [Figure 5d (right panel). In contrast, mammary epithelial cells had lower shape indices, reflecting a more compact and tightly packed arrangement [Figure. 5d (right panel)]. These differences were also in agreement with the PIV maps of the two wild-type tissues [Supplementary figure 2, Supplementary videos 4 and 6] where the mammary monolayers underwent a sharper drop in velocities and therefore kinetic arrest, compared to the bronchial epithelia. Bronchial epithelial cells also exhibited weaker cell-cell adhesions, as indicated by reduced E-cadherin staining intensity at cell junctions compared to mammary epithelial cells [Figure. 5c (bottom panel), e]. These weaker adhesions likely contributed to increased cellular motility and a diminished ability to constrain mutant clusters, in contrast to the more stable, compacted clusters observed in mammary epithelium. Together, these findings demonstrate that variations in mechanical properties-including cell shape, actin organization, and adhesion strength-govern the differential responses of mammary and bronchial epithelia to oncogenic mutants and may in turn, influence tumor growth and progression.

## Discussion

A central challenge in cancer biology is understanding why some oncogenic mutations are more potent in certain cancers. Large-scale cancer genomics has revealed that many driver mutations are shared across epithelial cancers, yet tumor incidence, progression rates, and invasive potential vary widely between organs ^[38, 39]^. These observations have traditionally been explained through tissue-specific differences in signaling pathways, differentiation states, and mutational landscapes. However, such explanations are incomplete, particularly in the context of early tumorigenesis, where oncogenic mutations are often detected in histologically normal tissues without leading to overt cancer ^[38]^. This raises a fundamental question: **what prevents or permits oncogenic cells from expanding at the earliest stages of cancer initiation?**

The present study addresses this gap by proposing interfacial mechanics between wild-type and nascently transformed cells, as a central organizing principle in early cancer initiation. Rather than focusing oncogenic cells in isolation, our work frames tumor initiation as an emergent outcome of mutant–wild-type interactions within an epithelial collective. From this perspective, the critical determinant of oncogenic fate is not only the presence of a mutation, but how the host tissue mechanically responds to it. This conceptual shift aligns with growing recognition that epithelial tissues behave as active materials, whose collective mechanical state can either suppress or amplify local perturbations. These insights shed light on the differential susceptibility of epithelial tissues to oncogenic transformation and uncover a mechanical basis for tissue-specific oncogenesis. By demonstrating how interfacial tension at the mutant-wild-type boundary governs whether oncogenic cells are restrained or overgrow, our work shifts the focus from purely molecular pathways to the biophysical constraints that regulate tumor initiation.

While previous studies have emphasized the contextual nature of tumorigenesis ^[2-4]^ and highlighted the critical role of interactions between healthy and oncogenic cells during epithelial cancer initiation ^[14, 22, 40]^, our work integrates and extends these perspectives by demonstrating how interfacial tension at the mutant-wild-type boundary determines whether transformed cells remain constrained or undergo unchecked expansion.

Importantly, while classical frameworks based on interfacial tension have often been interpreted through the lens of the Differential Interfacial Tension Hypothesis (DITH) ^[31,32]^ which attributes segregation to global differences in the contractility and adhesive properties of two tissues displaying different intrinsic tensions, the results of the present work support a different scenario, where what counts is the difference in the heterotypic interfacial tension, i.e, the tension along the boundary of the wild-type and mutant populations. We demonstrate that this heterotypic interfacial tension is a key determinant of mutant cluster morphology and dynamics. This is also consistent with recent theoretical work showing that sharp interfaces can emerge from interfacial mechanics alone ^[41]^. When interfacial tension between wild-type and mutant populations was positive, mutant clusters shrunk into circular aggregates and stayed constrained. In contrast, a negative interfacial tension resulted in irregular, protrusive clusters that failed to compact and kept growing. In line with this, we also showed that disrupting the interfacial actomyosin belt around the mutant clusters in the mammary epithelia, disrupted the restraint on them, preventing their demixing and compaction and shifting cluster behavior to a more unjammed state, similar to the ones in the bronchial system. These findings underscore the role of interfacial cellular mechanics in shaping oncogenic cell fate, with mammary epithelium imposing physical constraints that limit mutant expansion, while bronchial epithelium lacks such mechanical barriers, permitting unchecked growth.

By integrating mechanical constraints into models of cancer initiation, our study provides a framework for understanding how different epithelial tissues resist or permit tumorigenesis. However, while interfacial tension emerges as a key regulator of oncogenic segregation, the molecular pathways driving these mechanical differences remain to be elucidated. Future studies should investigate how targeting specific signaling networks or cytoskeletal components influences interfacial tension and whether modulating these factors could provide therapeutic benefits. Additionally, extending this framework to other oncogenes and tissue types could further elucidate the mechanical principles governing tumor initiation and progression.

In conclusion, this work supports a growing view of cancer initiation as a mechanically regulated, tissue-dependent process rather than a purely mutation-driven event. By positioning interfacial mechanics between oncogenic mutant cells and their wild-type counterparts, alongside genetic and biochemical factors, our findings contribute to an emerging physical understanding of cancer, in which tissue mechanics play an active role in shaping oncogenic fate. Such a framework not only helps explain tissue-specific cancer susceptibility but also suggests that reinforcing mechanical barriers within epithelia may represent an underexplored strategy for limiting early tumor progression.

## Materials and methods

### Cell culture

Non-transformed human mammary epithelial cell line MCF10A were maintained in complete medium composed of phenol-free Dulbecco’s modified Eagle’s medium (DMEM–F12, Gibco) supplemented with 5% charcoal-stripped horse serum (Gibco), 10 U ml^−1^ penicillin and 10 μg ml^−1^ streptomycin (Pen-Strep, Invitrogen), epidermal growth factor (20 ng/ml; PeproTech), hydrocortisone (0.5 mg/ml; Sigma-Aldrich), cholera toxin (100 ng/ml; Sigma-Aldrich), and insulin (10 μg/ml; Sigma-Aldrich) at 37°C in a humidified incubator with 5% CO2.

Non-transformed human bronchial epithelium epithelial cell line BEAS2B were cultured in Dulbecco’s modified Eagle’s medium (DMEM, Gibco) supplemented with GlutaMax (Gibco) and 5% fetal bovine serum (FBS, Gibco) along with 10 U ml^−1^ penicillin and 10 μg ml^−1^ streptomycin (Pen-Strep, Invitrogen). Cells were maintained at 37 °C and 5% CO_2_ unless mentioned otherwise.

### Cell seeding

10^5^ Cells were seeded on fibronectin-coated (10 μg ml^-1^) glass bottom dishes (35 mm Cellvis) and transfected once they reached 80% confluency (for singlets) and 60% confluency to get transfected groups.

### Transfection

Cells were transiently transfected with the Hras^V12^-GFP expressing plasmid DNA (Addgene #18780) using Lipofectamine 3000 (Invitrogen), following the manufacturer’s protocol. Post-transfection, confluent monolayers were either fixed and immuno-stained or imaged directly.

### Blebbistatin addition

Blebbistatin was diluted to 5 μM final solution and added 48h post-transfection. Samples were washed after 24 hours and fixed right away.

### Immunofluorescence

Cells were fixed with 4% paraformaldehyde (ThermoFisher) diluted in 1X Phosphate buffered saline (PBS, Sigma-Aldrich) for 15 minutes at room temperature. After washing away the fixative with 1X PBS, cells were permeabilized with 0.3% TritonX-100 (Sigma-Aldrich) in 1X PBS. Nonspecific antibody binding was blocked by incubating the samples in blocking solution (2% Bovine Serum Albumin (BSA Sigma-Aldrich) and 2% FBS in PBS) at room temperature for 45 minutes. Further, cells were incubated with primary antibody prepared in blocking/staining solution overnight at 4°C. Post primary antibodies incubation, cells were washed thrice with 1X PBS for 5 minutes per wash and incubated with secondary antibodies conjugated to Alexa fluor 555, Alexa fluor 594 conjugated phalloidin and 4′,6-diamidino-2-phenylindole (DAPI) to mark the cell nucleus, prepared in blocking/staining solution, for 2 hours at room temperature. Finally, the samples were washed four times with 1X PBS for 5 minutes per wash before proceeding to microscopy.

### Microscopy

Fluorescence images were acquired using 20X objective and 63x oil-immersion objective of a Zeiss Axio Observer 7 Inverted Microscope with a scientific-grade sCMOS camera by Iberoptics. For live imaging, samples were set up on a stage-top incubator and maintained at 37°C and 5% CO2 throughout imaging.

### Image Analysis

Image analysis for this study was performed using Fiji ^[42]^ except bayesian force inference and particle image velocimetry (PIV) analysis, which was performed in MATLAB (MathWorks). Cellpose ^[43]^ was used to segment cells to get cell count, shape indices and actin intensitites for each cell while Tissue Analyzer ^[44]^ was used to segment cells for force inference.

### Quantification of cluster eccentricity and segregation index

The term ‘eccentricity’ expresses a size reduction in the contact with neighboring cells. It was quantified using the formula 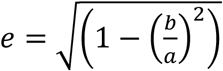, (where a= the length of the major axes and b= the length of the minor axes of the clusters), with a value closer to 0 indicating a more circular shape. Eccentricity quantifications were performed on fixed and live tissues using outlines that were manually drawn and measured using Fiji.

The segregation index SI, defined as the average ratio of homotypic and all cell neighbors as done previously ^[45]^, quantifies the demixing degree. To quantify the SI, cells of each type were manually counted, and finally, to generate the plots, the SI was averaged over all GFP-labelled cells in one frame.

### Bayesian Force Inference

Bayesian force inference was done as described previously ^[26]^ to determine the relative cellular pressures and cell-cell edge or line tension.

### Particle Image Velocimetry (PIV)

Particle Image Velocimetry (PIV) was done using the PIV Lab package on MATLAB ^[46]^. FFT window deformation (direct Fourier transform correlation with multiple passes and deforming windows) algorithm was used. Velocity maps were obtained by a 3-pass interrogation window of 128 × 128 pixels that halved after each pass. Tangential velocities were obtained by drawing concentric circles radially outward from the center to a point at the boundary of the mutant clusters and integrated over the entire circle using a custom python code. The output is the tangential velocity as a function of the circle.

### Statistical analysis

Statistical analyses were carried out in GraphPad Prism 10. Statistical significance was calculated by Unpaired t-test with Welch’s correction. p-values greater than 0.05 were considered to be statistically not significant. Each plot has data pooled from multiple frames of one out of three independent experiments. Violin plots have median shown as a bold dashed line and the first and third quartiles shown as thin dashed lines. Box and whisker plots have whiskers extending from the minimum to the maximum values, showing the full range of the dataset including outliers. Scatter-bar plots were displayed as mean ± s.e.m. No statistical methods were used to set the sample size. Quantification was done using data from at least three independent biological replicates. For analysis involving live-imaging experiments, data were collected from three independent experiments. All the experiments with representative images were repeated at least three times.

### Vertex Model

A bi-disperse model is used to demonstrate the role of interfacial tension between two types of cells, specifically the HRas^V12^ mutant and wild-type cells.

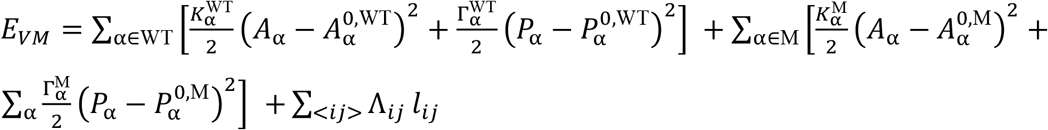

The first two sums are included in the standard vertex model, whereas the last term is introduced in our model to account for interfacial interactions between cells. Specifically, Λ_*ij*_ = 0 for interfaces between same cell types, and Λ_*ij*_ = Λ between different cell types. Using a line tension term like Λ*l*_*ij*_ in the vertex model energy, where the *l*_*ij*_ is the length of an interface edge, we can obtain both segregation and spreading dynamics just from energy minimization. This term has been described as heterotypic line tension (HLT)^[47]^. Due to energy minimization, heterotypic interface length is minimized for Λ > 0 and maximized for Λ < 0, leading to segregation of the two cell types in the former case and complete mixing of them in the latter case.

The mutant cluster in the experiments with MCF10A cells showed an actin belt forming along the cluster interface, with the cells along the interface being more elongated than normal. This actin ring is important for the compaction of mutant clusters in the presence of wild-type cells in the epithelial monolayer. An isotropic interfacial tension, such as the Λl_ij_ term in our model cannot capture such anisotropic behaviour. Recently, it has been proposed that such anisotropic behaviour can arise from the anisotropic distribution of the cytoskeletal filaments inside the cell^[40]^. The resulting active interfacial tension, called the shape-tension coupling, can generate elongated shapes as observed in the experiments. The elongated cells in the experiments indeed have anisotropic distributions of the stress fibres. Hence, we included the effect of shape-tension in our model. The total interfacial tension is Λ_*ij*_ + *γ*_*ij*_(*t*; Π_0_), where Λ_*ij*_ is a time-independent, passive, isotropic tension and *γ*_*ij*_ is a time-dependent active tension that depends on the shape of the cells sharing the interface and requires a threshold stress Π_0_ to be nonzero. The details of the shape-tension coupling is described in the appendix. We find that a nonzero activation stress Π_0_ is necessary to reproduce experimentally observed results.

## Appendix

### Vertex Model Simulation

To understand the cause and the dynamics of interaction between mutant and wild-type population in different epithelial monolayers during cancer initiation, we used a cell-based model of epithelial tissues called the vertex model. A monolayer in the vertex model has total energy given by:

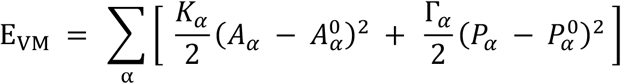

In the above energy function, the constant parameters *K*_*α*_ is the area modulus which gives the area elasticity of each cell and Γ_*α*_ is the perimeter modulus which give the elasticity due to the actomyosin cortex. The gradient of this energy function gives the forces on each vertex of the cell, 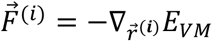, where 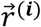 is the position of the *ith* vertex. The dynamics of each vertex position is given by the overdamped equation of motion:

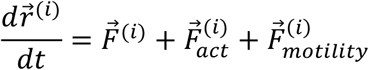

where 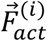 is any active force which might drive the system out of equilibrium. 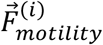is a force which propels individual cells by 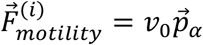, where 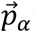 is the polarity of a given cell given by 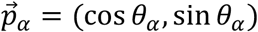, where this angle *θ*_*α*_ performs rotational diffusion:

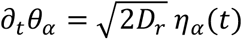

where *η*_*α*_ (*t*) is a Gaussian white noise with zero mean and correlation ⟨*η*_*α*_ (*t*) *η*_*α*_′(*t*′)⟩ = *δ*(*t* − *t*′) *δ*_*α α*_′.

To capture the dynamics of cell segregation between two different types of cells in a monolayer of normal and oncogenic cells, we can use a bi-disperse vertex model with an interfacial line tension between the two types of cells - called mutant and wild type can be used.

The director of each cell corresponds to the polarization direction of the cell, which can be obtained as the larger eigenvector of its shape tensor, 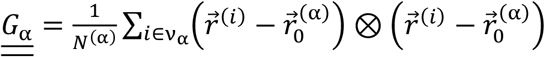, where the sum is over *N*^(*α*)^ vertices of the cell *α*, and 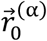 is the position of the cell’s geometric center. The energy function for the vertex model used is given below:

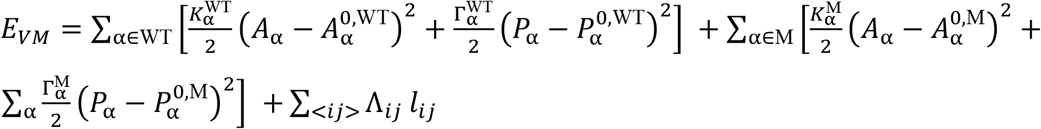

Here, *M* denotes mutant cells and *WT* denotes wild-type cells. The index *α* denotes cells, and the indices *i, j* denote vertices and < *ij* > denotes an edge between vertices *i* and *j*. The parameters 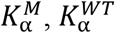 are the area moduli, 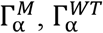 are the perimeter moduli of the mutant and wild type respectively, and the Λ_*ij*_ denotes the interfacial line tension. This interfacial line tension is only along junctions connecting two vertices and depends on the cell type on either side of the junction. Since the system we are considering is bi-disperse, the interfacial line tension can have three values, depending on the identities of cells on either side of a given edge < *ij* >:

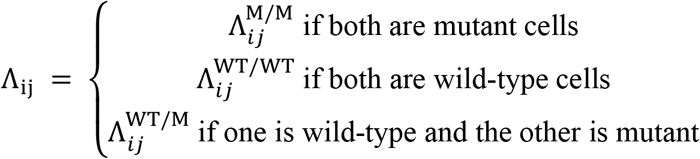

### Shape-tension coupling

In addition to this passive interfacial tension, we can also have an active contribution coming from the shape tension coupling. The shape-tension coupling is mathematically the same as the passive line tension force. The total force is given by:

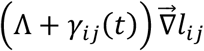

*γ*_*ij*_(*t*) depends on the relative orientation of the cell’s director 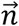 and the edge along which the tension acts, 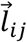 Specifically, it has a relaxational dynamics of the form:

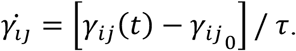

Here 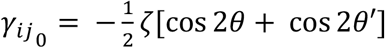 couples the tension along an edge to the shape of the cells on either side of the edge [Appendix Fig 1].

For ζ > 0, this tension is extensile and leads to the elongation of cells along the boundary. This happens because for the edges which have 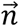 and 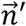 aligned with the edge vector (*θ* or *θ*′ = 0, π, 2π), the shape-tension coupling contributes a negative line tension, which favours this alignment. The directors can be obtained from the larger eigenvector of the shape tensor of the cells as described before. When the edge vectors are not aligned with the edge vector, the contribution is positive, and that edge starts to shrink. Eventually, the interface consists of cells whose directors are always aligned with the interface. Energetically, it counteracts interfaces with Λ > 0, leading to the formation of elongated cells and tangential flows along the interface. Importantly, we find that cell elongation correlates with the formation of actin belt, leading us to postulate that the shape-tension coupling is activated above a threshold pressure, Π_’_, where the cell pressure is given by 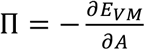, When Λ < 0, such as in the BEAS2B cells, we do not observe strong stress-fibre expression. Hence, we believe that it is not activated in those cells.

### Simulation Setup

The vertex model energy can be non-dimensionalised by setting a length scale set by 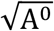. In our simulation we have taken

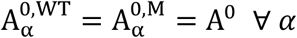

and

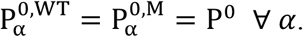

Also,

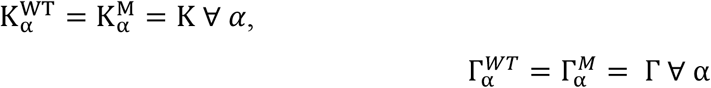

and

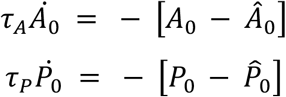

The non-dimensionalization has been done, which defines the normalized parameters 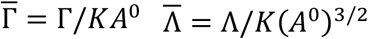 . (In further discussions, the normalized parameters will be represented by the symbols Γ and Λ.)

The monolayer modelled by the vertex model can be described by a single number, the shape index, defined by 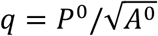 . Here too, in principle we could have different shape indices for the mutant and wild type cells, but we take them to be the same. Unless otherwise mentioned, we take Λ^*M/M*^ = Λ^*WT/WT*^ = 0 and Λ^*WT/M*^ = Λ.

To simulate the elongation of cells along the interface, ζ > 0 is used for an active extensile contribution to the interfacial tension.

A box containing 20 × 20 cells has been used with mutant cells placed within a region (non-circular) from the center of the box. The vertices are arranged in a hexagonal lattice with random displacements added to the position of each vertex. Periodic boundary conditions were used to simulate a confluent monolayer.

### Parameters

*Parameters:*
|  |  |
| --- | --- |
| $K$ | 1 |
| $\Gamma$ | 0.1 |
| $A^0$ | 1 |
| $P^0$ | 4.0 |
| $l_{min}$ for T1 transitions | 0.04 |
| $\Delta t$ for time stepping | 0.01 |
| $v_0$ for motility | 0.05 |
| $D_r$ for rotational diffusion | 0.5 |
| $\zeta$ for shape-tension coupling | 0.6 |

To prevent overlaps, a node switch operation is also added to the simulation, which changes the topology of the network ^[21]^.

### Extrusion Probability Calculation

Simulations with just a single mutant cell were run for a range of heterotypic interfacial line tension values (Λ = 0, 0.1, 0.4, 0.8, 1.2, 1.6) with shape tension coupling. The simulation was run till the area of the mutant cell fell below a threshold area = 0.1, after which we consider the mutant cell to be extruded. 9 different random initial seeds were run and analysed. Each seed gives a binary result – either extruded or not. This was used to calculate the extrusion probability.

### Need for Shape-Tension Coupling – Elongation Along Interface

Shape-tension coupling has been used here in accordance with the experimental observation that the cells at the interface are aligned and elongated along the interface [Fig. 2h]. Below, difference between shape indices of cells at the interface and away from the boundary is plotted versus the interfacial tension in the case of no shape-tension coupling [Appendix Fig 2]. The red dashed line represents the experimental value of the shape index difference. The blue line is the shape index difference between two randomly chosen groups of cells (half of the total number of cells in each group is taken). At zero line-tension, the difference in shape index between interface cells and cells away from the interface is same as that between randomly chosen groups of cells, which is expected since there should be no interface at zero line-tension. The no shape-tension data presented here are averaged over 19 seeds. Although the results without shape-tension coupling reaches experimental values at high enough interfacial tension [Appendix Fig 2], a closer inspection of the simulation results show that the cells are just squeezed and are aligned perpendicular to the interface, which is contrary to what is seen in experiments [Fig. 2h].

Calculating the average of the absolute value of the dot product of the nematic director and the interface edge for simulations with and without shape-tension coupling [Appendix Fig 3] clearly shows that with shape-tension coupling, the cells align and elongate along the interface as is seen in experiment, given by an interface dot product value > 0.5 at high enough line-tension values. Further, shape-tension coupling or biased edge tension has been used before to model for cell elongation during embryo elongation ^[48]^ and here we use it as an active line-tension force, which elongates cells along the interface, in addition to the interfacial tension which is passive.

### Vertex Model with Mechanochemical Feedback

Although the difference in actin between mutant and wild-type have not been incorporated in the model presented in the manuscript, we could see how the actin levels change in response to the interfacial tension formed between the mutant and wild-type cells by adding a mechanochemical feedback in the model. As given in^[35]^, we can model the behaviour of tissues using a vertex model with mechanochemical feedback. It has been shown that incorporating these feedback loops in the vertex model captures the biologically realistic features of epithelial tissues seen in experiment^[35]^. In particular, to model the actin distribution in monolayers with mutants and wild-type, we can use a vertex model with interfacial tension and MCFL-I^[35]^. When we incorporate MCFL-I in our model, we can observe how the normalized contractility 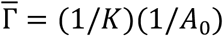, and normalized line tension 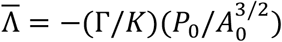 changes with different line tensions. The normalized contractility is associated with the bulk actin or the bulk compressibility, and the normalized line tension is associated with the junctional actin of the cells. The equations for MCFL-I can be written as:

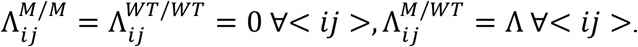

Thus, with MCFLs, the vertex model does not have fixed A_0_ and P_0_. The cells dynamically change these parameters depending on the vertex model dynamics. The constitutive relations for the 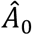 and 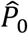 are given below^[35]^:

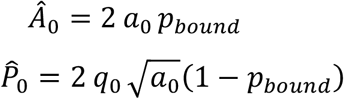

Here, p_bound_ = 1/(1 + (A/Â)^2^), which is the probability of myosin binding and Â is a model parameter which depends on the dissociation constant of myosin^[35]^. We consider Â to the be the same for both mutant and wild-type (Â = 1.3). A positive heterotypic interfacial tension can lead to compression (decrease in area) of the mutants whereas a negative heterotypic interfacial tension can lead to relaxation of cell area of the mutants. This will lead to different A_0_ and P_0_ across the monolayer, and thus different 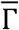 and 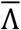, which provides an understanding of the different actin levels.

For positive heterotypic interfacial tension, the mutants are compressed which leads to decrease 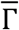 and an increase in 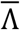 (Figure 1). In the figures below, we plot the 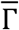 as the bulk actin and the 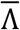 as the junctional actin. Since our model has an active line tension, the normalized line tension of an edge will also have a contribution from Λ, thus 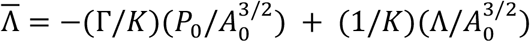. The spatial distribution of actin in the mutant and wild-type seen in experiments matches well with the snapshot from simulations, where the bulk actin is low within the cluster as compared to the wild-type [Appendix Fig 4]. For negative heterotypic interfacial tension too, we see that the actin levels are not very different between the mutant cluster and the wild-type [Appendix Fig 5].

## Author contributions

M.V conceived the project. M.V. and A.D. designed experiments. A.D performed all experiments except the staining on BEAS2B cells, which were performed by A.A. Theoretical model was contributed by P.D and S.S. Analysis and interpretation of experimental data was done by A.D, T.C, T.T, A.A, S.M, A.R and M.V. A.D and M.V developed and wrote the manuscript with help from P.D and S.S. All authors read, discussed and commented on the manuscript.

## Acknowledgements

We thank Sriram R. Ramaswamy, and Tamal Das for critical discussions and suggestions. M.V. is a partner group leader of the Max Planck Society (MPG), Germany, which has supported part of this work. This work is also supported by the Infosys foundation, Anusandhan National Research Foundation-previously called the Science and Engineering Research Board (project number: SERB SRG/2022/000534), and Indo German Science and Technology Centre (IGSTC WISER scheme). SS acknowledges funding from IISc, Axis Bank Center for Mathematics and Computing, and a startup grant from SERB-DST (SRG/2022/000163). We also acknowledge intramural funds at IISc Bangalore for providing support towards equipment and facilities and for salaries/fellowships of the author

## Supplementary

**Supplementary Figure 1:**
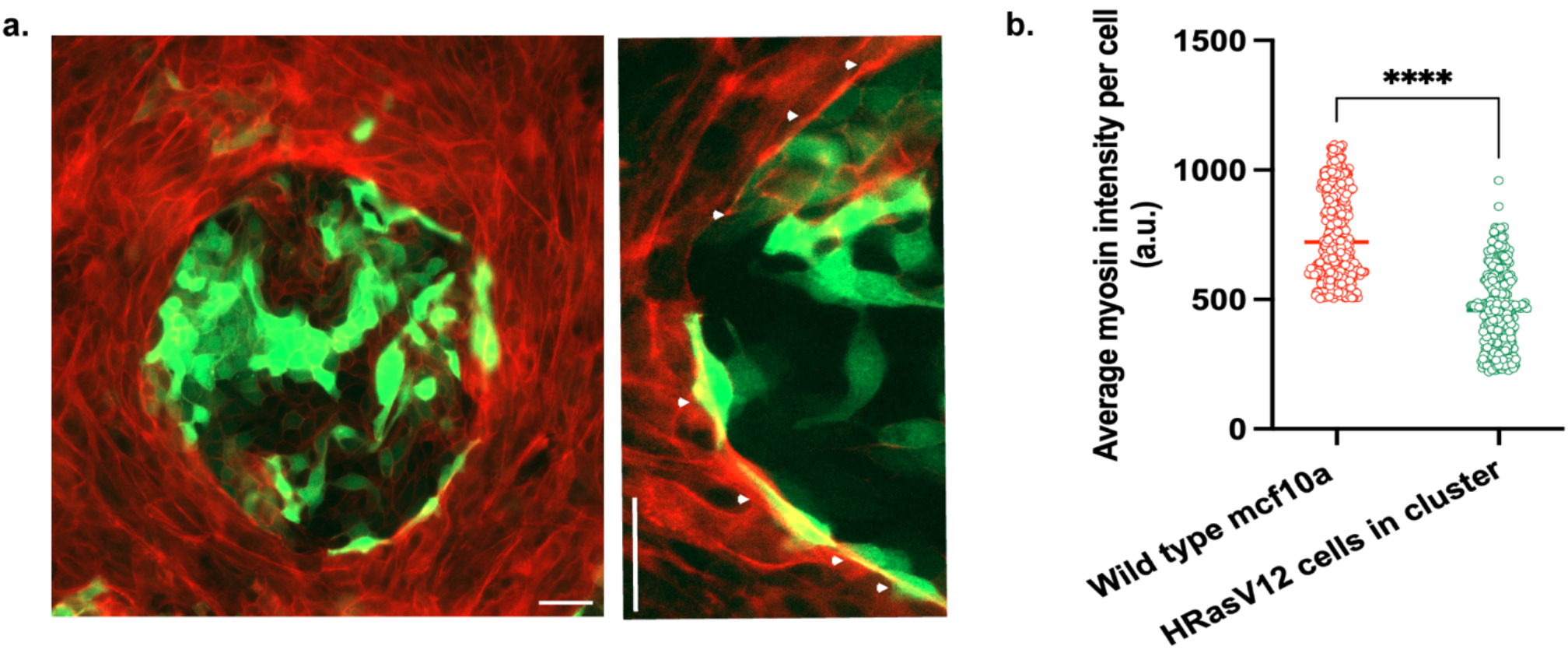
Myosin belt and per cell myosin intensities. (a) Images of ^HRasV12^ cluster in mammary epithelia stained for phosphomyosin (left panel) and phosphomyosin belt around the cluster marked with white arrowheads (right panel) (b) Plot showing lower phosphomyosin levels inside oncogenic clusters in MCF10A Representative data are plotted from one of three independent experiments with the median shown as a bold line. Statistical significance was calculated using Unpaired t-test with Welch’s correction. *Scale bars= 50 μm*.

**Supplementary Figure 2:**
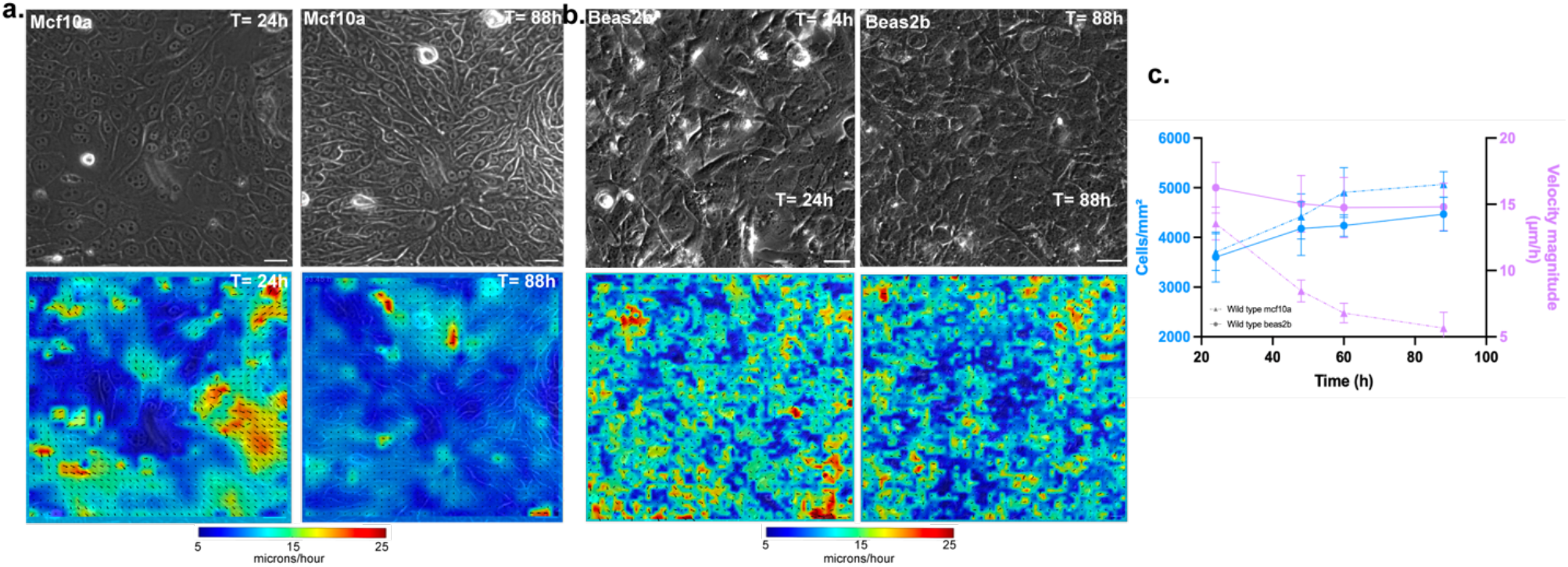
PIV maps of wild-type mammary and bronchial epithelial monolayers with increasing cell density. (a) Representative time-lapse images of wild-type (control) mammary epithelium at two different time points, with corresponding PIV velocity maps showing a reduction in overall velocity as cell density increases (b) Representative time-lapse images of wild-type (control) bronchial epithelium at two different time points, with corresponding PIV velocity maps demonstrating a similar reduction in velocity with increasing cell density (c) Quantification of velocity reduction in wild-type mammary and bronchial epithelia as a function of cell density, highlighting tissue-specific differences in collective cell dynamics. All data are represented as mean±sem plotted from different clusters from one representative experiment. *Scale bars= 50 μm*.

**Supplementary Figure 3:**
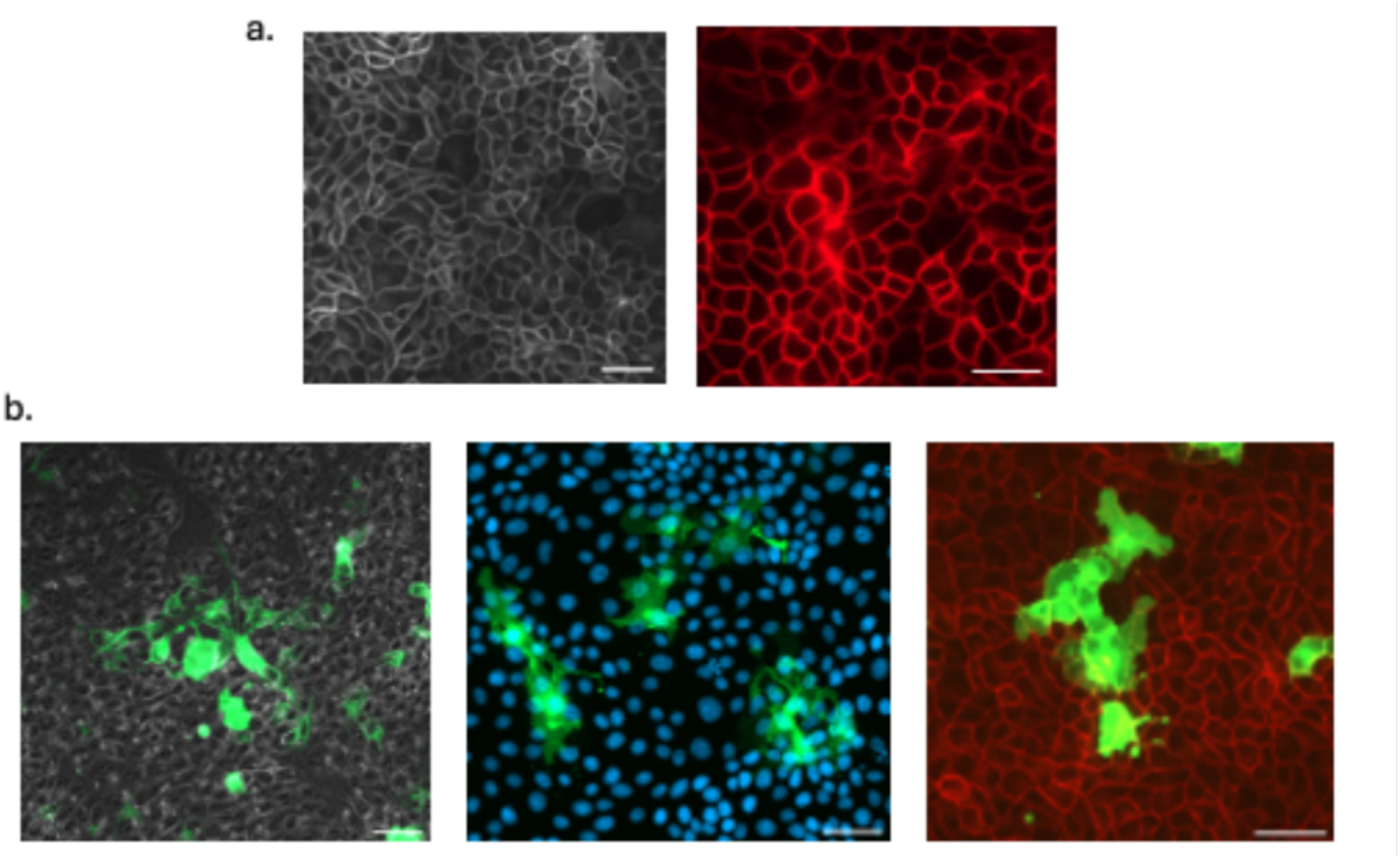
HRas^V12^ clusters show invasive phenotype in MDCK monolayer. (a) Representative images of wild-type MDCK monolayers stained for F-actin (left) and E-cadherin (right), showing intact cell-cell junctions. (b) HRas^V12^ clusters expand, forming long protrusions over time. *Scale bars= 50 μm*.

**Supplementary Figure 4:**
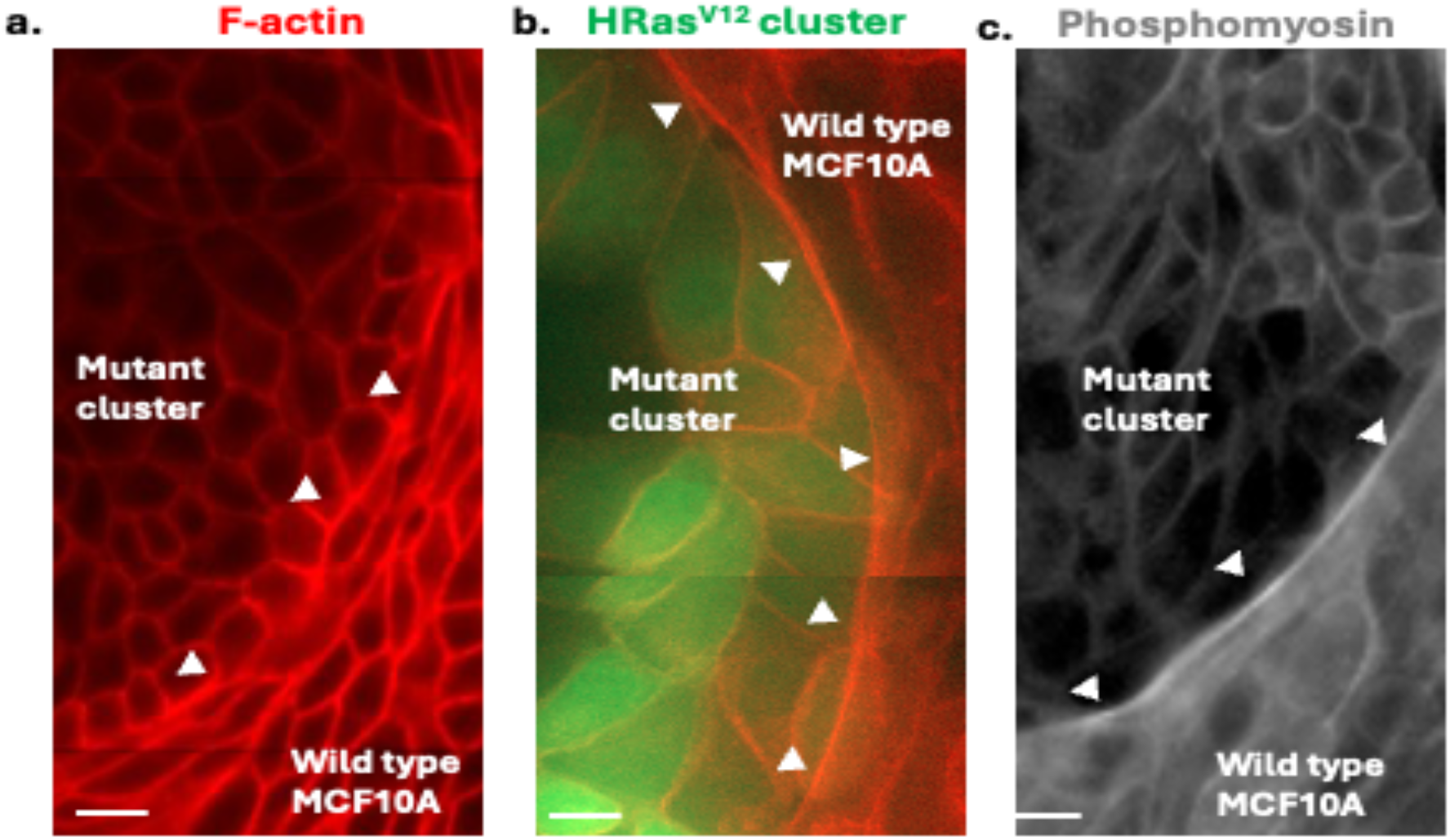
Actomyosin enrichment at the interface of HRas^V12^ clusters and wild type MCF10A. (a) Images of HRas^V12^ cluster in mammary epithelia stained for F-actin, showing higher F-actin deposition at heterotypic interfaces (b) Representative image of HRas^V12^ cluster in mammary epithelia stained for Phosphomyosin, showing higher myosin deposition at the heterotypic interface. *Scale bars= 10 μm*.

**Supplementary Figure 5a:**
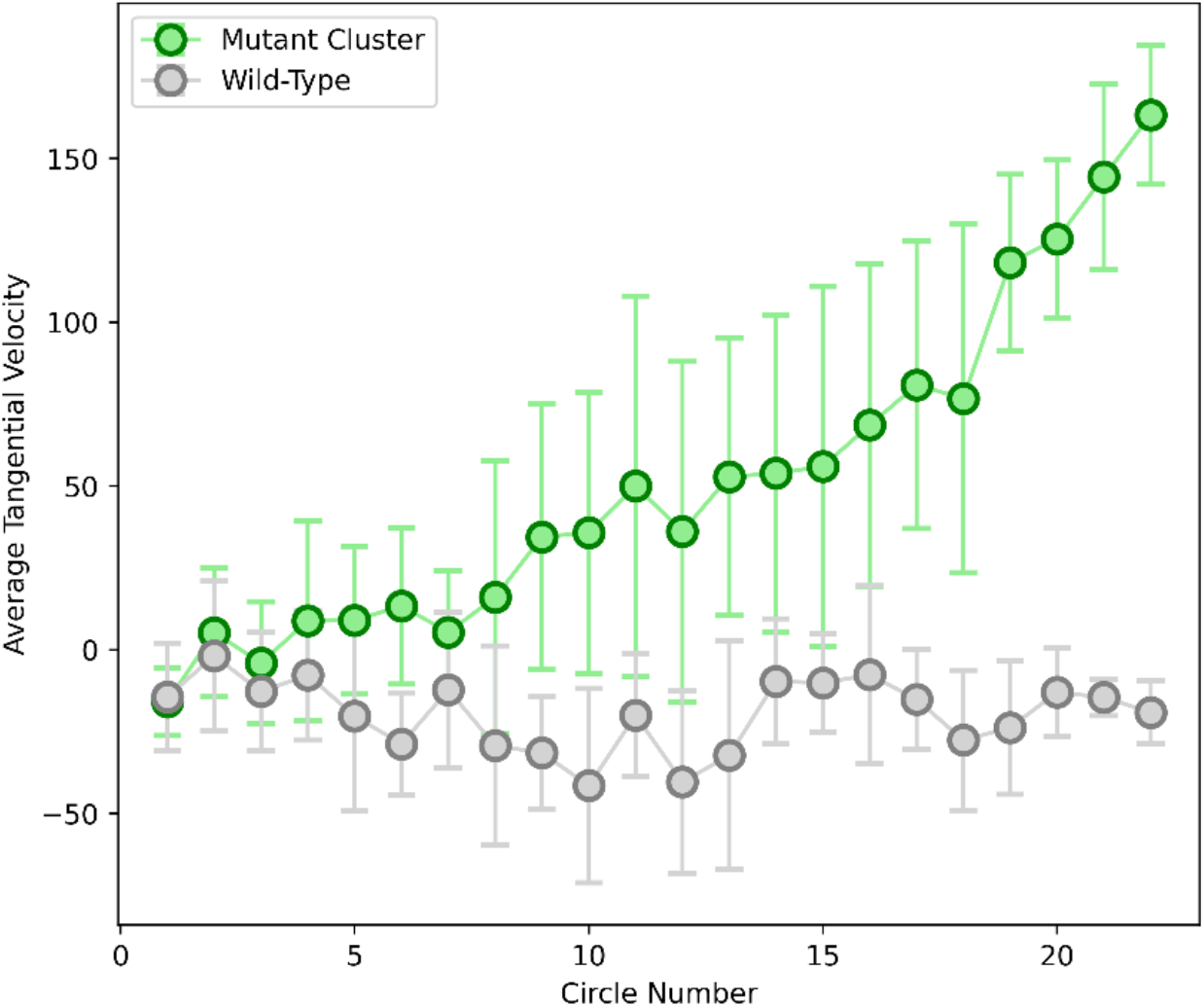
Normalized tangential velocity loop integral plotted vs. circle number (increasing radius). The integral has been normalized by the perimeter of the circle. The same trend as in the un-normalized analysis is seen. Average has been taken over 8 different wild-type and mutant cluster positions.

**Supplementary Figure 5b:**
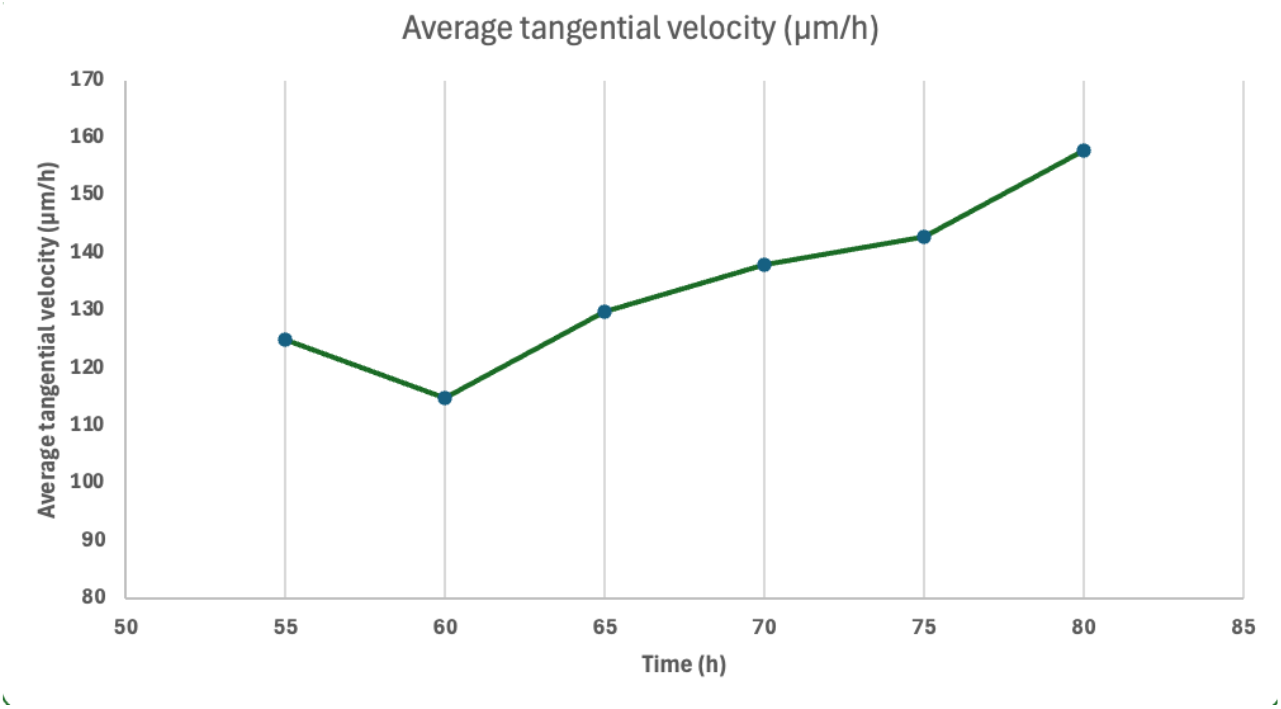
Normalized tangential velocity loop integral plotted for circle number 22 (cells at the interface of the mutant clusters) vs. time. The integral has been normalized by the perimeter of the circle. We see tangential velocity persists for over 24 hours, till the end of the time-lapse video, starting when the mutants have formed a mature compact, rounded, well defined cluster.

**Supplementary Figure 6:**
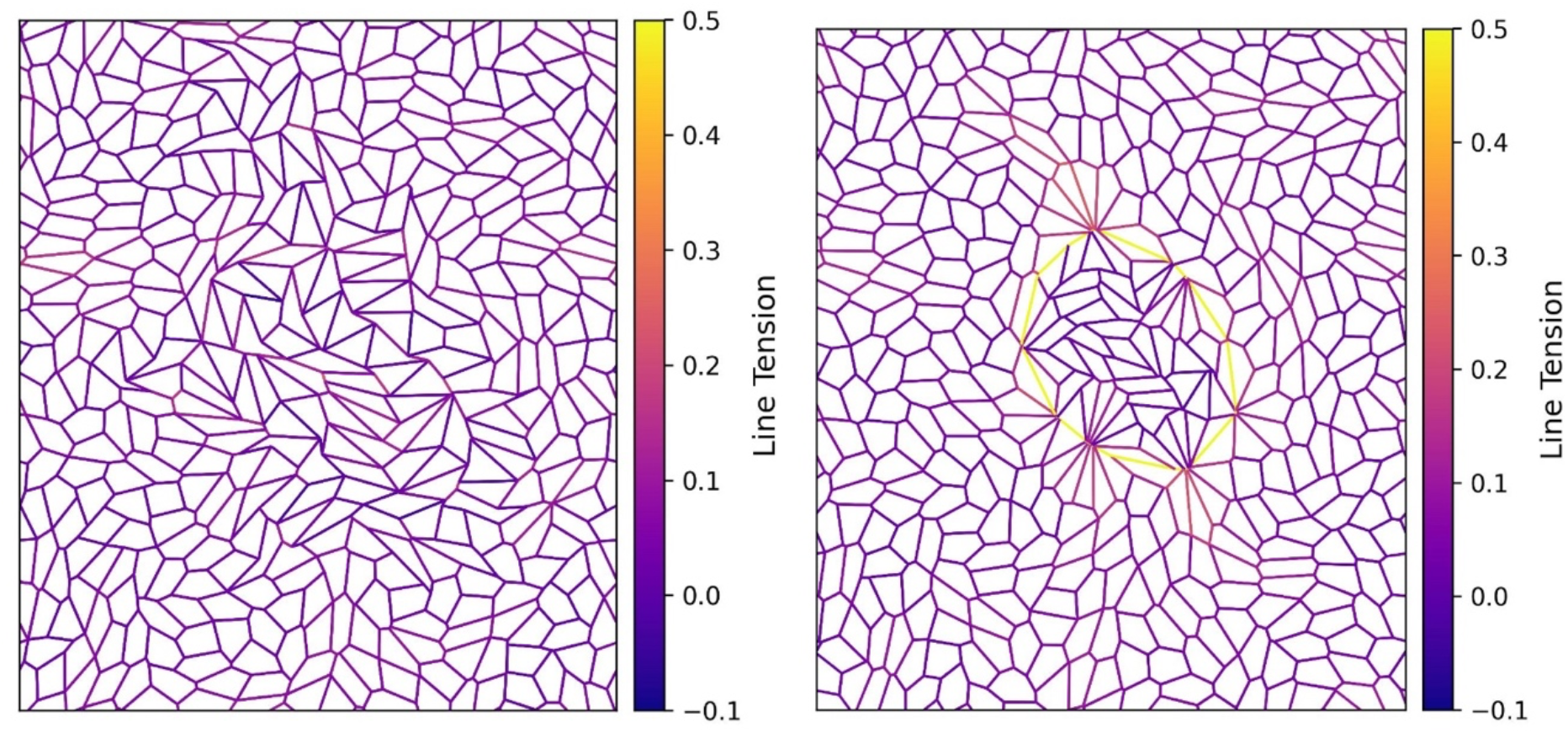
Line tensions from simulation for (*left panel*) Λ = −0.2, and (*right panel*) Λ = 1.6.

### Videos

**Supplementary video 1:** An HRas^V12^ singlet is extruded from MCF10A monolayer but persists in BEAS2B monolayer, multiply and form long protrusion. *Scale bar= 20 μm*.

**Supplementary video 2:** A group of HRas^V12^ transfected cells gradually gets confined into a compact cluster in MCF10A monolayer and forms a smooth interface with the wild-type population, while a group of HRas^V12^ transfected cells gradually grows in size in BEAS2B monolayer, forming a jagged interface with the wild-type population along with protrusions. *Scale bar= 50 μm*.

**Supplementary video 3:** PIV video during formation of a representative HRas^V12^ cluster in MCF10A monolayer. *Scale bar= 50 μm*.

**Supplementary video 4:** PIV video showing the dynamics of a representative wild-type MCF10A (control) monolayer without any oncogenic mutants. *Scale bar= 50 μm*.

**Supplementary video 5:** PIV video showing the dynamics of a representative HRas^V12^ cluster in BEAS2B monolayer *Scale bar= 50 μm*.

**Supplementary video 6:** PIV video showing the dynamics of a representative wild-type BEAS2B (control) monolayer without any oncogenic mutants. *Scale bar= 50 μm*.

**Supplementary video 7:** Video showing output of the bi-disperse vertex model with cluster dynamics varying with interfacial tension (Λ).

---

**Appendix Fig 1:**
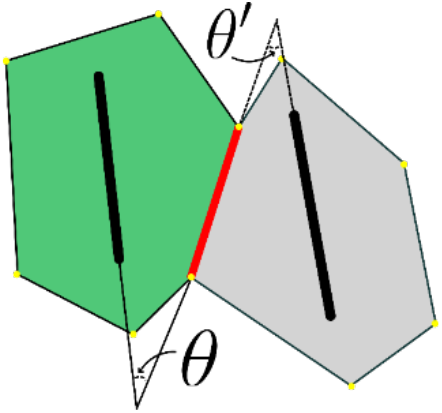
Illustration of shape-tension coupling at the mutant (green)–wild-type (gray) interface (red), where angles θ and θ’ represent the orientation of boundary cells relative to their shared edge.

**Appendix Fig. 2:**
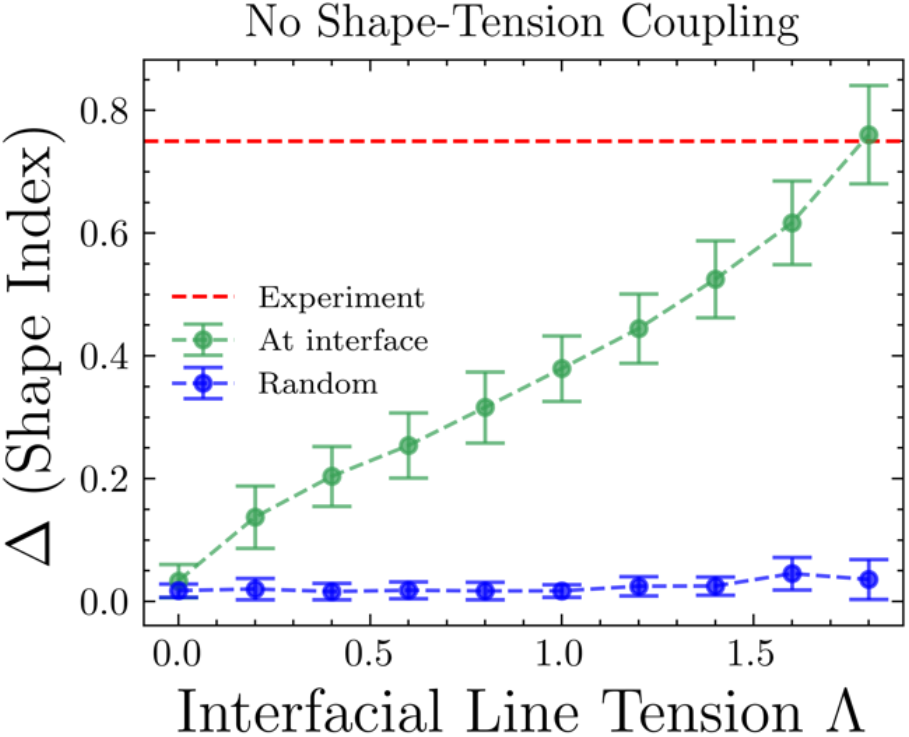
Shape indices versus the interfacial line tension for no shape-tension coupling.

**Apppendix Fig 3:**
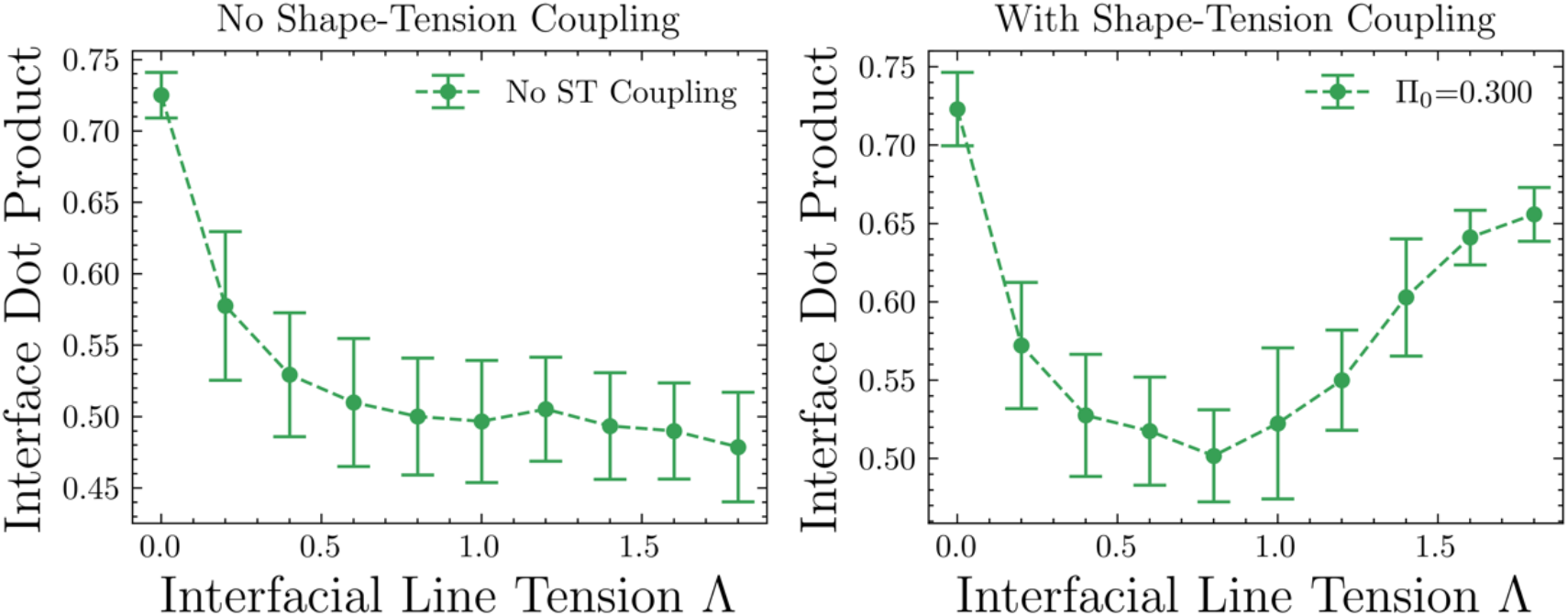
Interface dot product for different interfacial line tension with and without shape-tension coupling.

**Appendix Fig 4:**
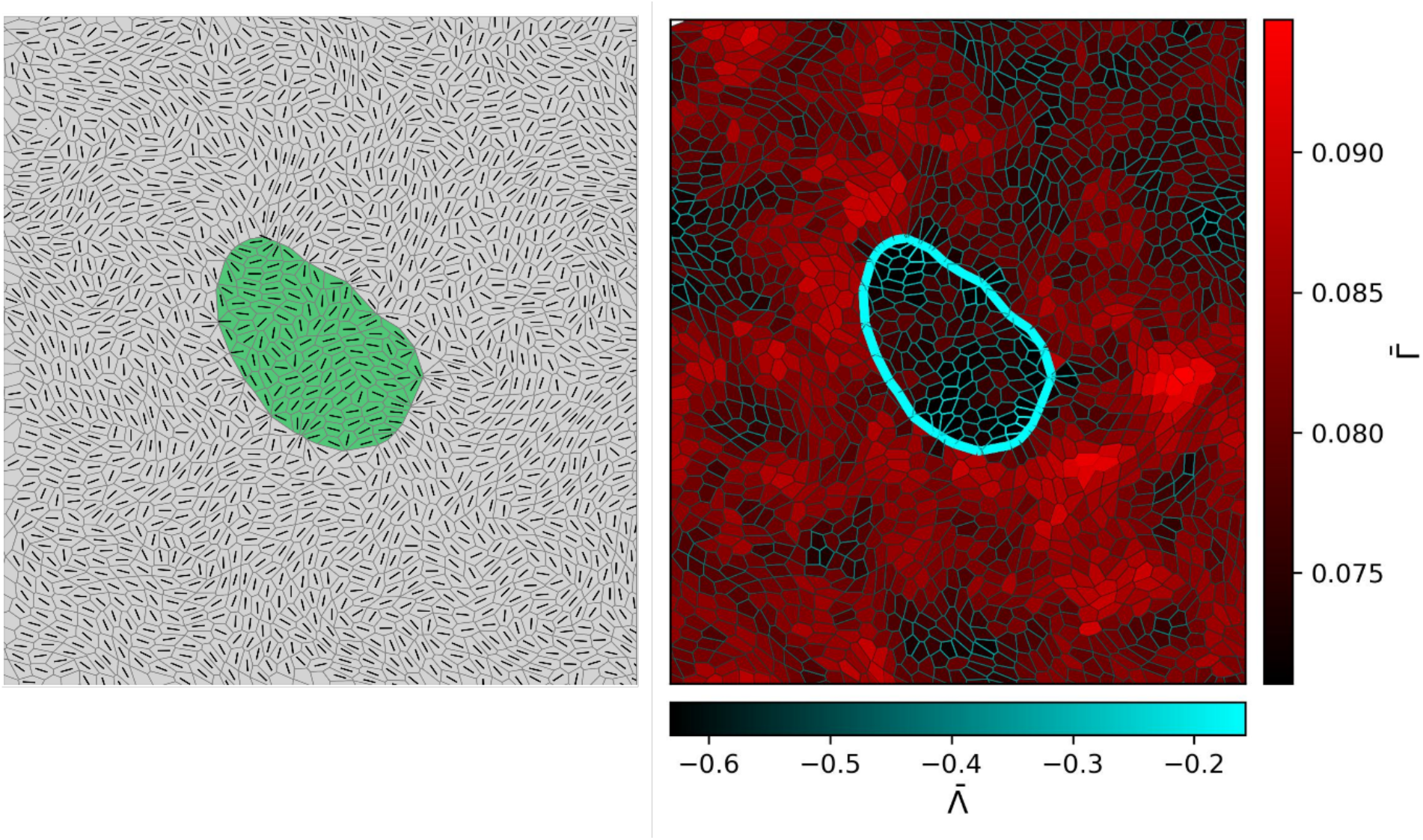
For positive differential interfacial tension (Λ >0), the mutant cluster has lower bulk actin level compared to the wild-type cells surrounding it.

**Appendix Fig 5:**
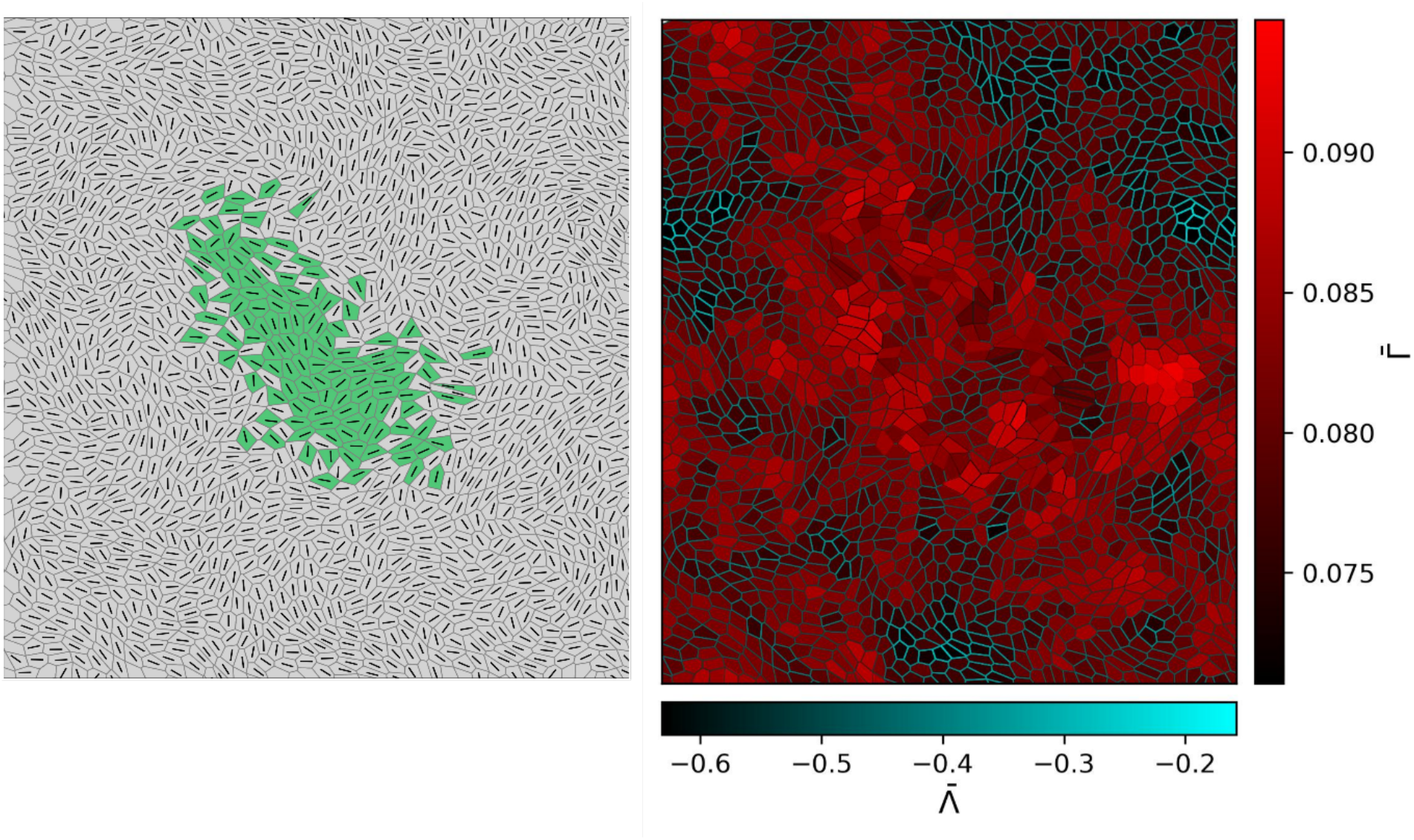
For negative differential interfacial tension (Λ <0), the mutant cluster does not have a very different actin level compared to the wild-type cells surrounding it.

## References

1. Hanahan D, Weinberg RA. 2000. The hallmarks of cancer. Cell 100:57–70. doi:10.1016/S0092-8674(00)81683-9.

2. Schneider G, Schmidt-Supprian M, Rad R, Saur D. 2017. Tissue-specific tumorigenesis: context matters. Nat Rev Cancer 17:239–253. doi:10.1038/nrc.2017.5.

3. Haigis KM, Cichowski K, Elledge SJ. 2019. Tissue-specificity in cancer: The rule, not the exception. Science 363:1150–1151. doi:10.1126/science.aaw3472.

4. Bianchi JJ, et al. 2020. Not all cancers are created equal: Tissue specificity in cancer genes and pathways. Curr Opin Cell Biol 63:135–143. doi:10.1016/j.ceb.2020.01.006.

5. Yamashita S, et al. 2018. Genetic and epigenetic alterations in normal tissues have differential impacts on cancer risk among tissues. Proc Natl Acad Sci U S A 115:1328– 1333. doi:10.1073/pnas.1717960115.

6. Beroukhim R, et al. 2010. The landscape of somatic copy-number alteration across human cancers. Nature 463:899–905. doi:10.1038/nature08822.

7. Zack TI, et al. 2013. Pan-cancer patterns of somatic copy number alteration. Nat Genet 45:1134–1140. doi:10.1038/ng.2760.

8. Davoli T, et al. 2013. Cumulative haploinsufficiency and triplosensitivity drive aneuploidy patterns and shape the cancer genome. Cell 155:948–962. doi:10.1016/j.cell.2013.10.011.

9. Salmon H, Remark R, Gnjatic S, Merad M. 2019. Host tissue determinants of tumour immunity. Nat Rev Cancer 19:215–227. doi:10.1038/s41568-019-0125-9.

10. Lehmann B, et al. 2017. Tumor location determines tissue-specific recruitment of tumorassociated macrophages and antibody-dependent immunotherapy response. Sci Immunol 2:eaah6413. doi:10.1126/sciimmunol.aah6413.

11. Kajita M, Fujita Y. 2015. EDAC: Epithelial defence against cancer—cell competition between normal and transformed epithelial cells in mammals. J Biochem 158:15–23. doi:10.1093/jb/mvv049.

12. Ohoka A, et al. 2015. EPLIN is a crucial regulator for extrusion of RasV12-transformed cells. J Cell Sci 128:781–789. doi:10.1242/jcs.159772.

13. Kajita M, et al. 2014. Filamin acts as a key regulator in epithelial defence against transformed cells. Nat Commun 5:4428. doi:10.1038/ncomms5428.

14. Hogan C, et al. 2009. Characterization of the interface between normal and transformed epithelial cells. Nat Cell Biol 11:460–467. doi:10.1038/ncb1853.

15. Pothapragada SP, et al. 2022. Matrix mechanics regulates epithelial defence against cancer by tuning dynamic localization of filamin. Nat Commun 13:218. doi:10.1038/s41467-02127640-w.

16. Sasaki A, et al. 2018. Obesity suppresses cell-competition-mediated apical elimination of RasV12-transformed cells from epithelial tissues. Cell Rep 23:974–982. doi:10.1016/j.celrep.2018.03.108.

17. Leung CT, Brugge JS. 2012. Outgrowth of single oncogene-expressing cells from suppressive epithelial environments. Nature 482:410–413. doi:10.1038/nature10826.

18. Alt S, Ganguly P, Salbreux G. 2017. Vertex models: from cell mechanics to tissue morphogenesis. Philos Trans R Soc Lond B Biol Sci 372:20150520. doi:10.1098/rstb.2015.0520.

19. Barton DL, et al. 2017. Active vertex model for cell-resolution description of epithelial tissue mechanics. PLoS Comput Biol 13:e1005569. doi:10.1371/journal.pcbi.1005569.

20. Fletcher AG, et al. 2014. Vertex models of epithelial morphogenesis. Biophys J 106:2291– 2304. doi:10.1016/j.bpj.2014.04.021.

21. Fletcher AG, Osborne JM, Maini PK, Gavaghan DJ. 2013. Implementing vertex dynamics models of cell populations in biology within a consistent computational framework. Prog Biophys Mol Biol 113:299–326. doi:10.1016/j.pbiomolbio.2013.09.003.

22. Gupta P, Kayal S, Tanimura N, Pothapragada SP, Senapati HK, Devendran P, Das T. 2024. Mechanical imbalance between normal and transformed cells drives epithelial homeostasis through cell competition. bioRxiv. doi:10.1101/2023.09.

23. Sinjab A, Han G, Wang L, Kadara H. 2020. Field carcinogenesis in cancer evolution: what the cell is going on? Cancer Res 80:4888–4891. doi:10.1158/0008-5472.CAN-20-2312.

24. Franklin WA, Gazdar AF, Haney J, Wistuba II, La Rosa FG, Kennedy T, Miller YE. 1997. Widely dispersed p53 mutation in respiratory epithelium. A novel mechanism for field carcinogenesis. J Clin Invest 100:2133–2137. doi:10.1172/JCI119748.

25. Curtius K, Wright NA, Graham TA. 2018. An evolutionary perspective on field cancerization. Nat Rev Cancer 18:19–32. doi:10.1038/nrc.2017.102.

26. Ishihara S, Sugimura K. 2012. Bayesian inference of force dynamics during morphogenesis. J Theor Biol 313:201–211. doi:10.1016/j.jtbi.2012.08.026.

27. Atia L, Bi D, Sharma Y, Mitchel JA, Gweon B, Koehler SA, Fredberg JJ. 2018. Geometric constraints during epithelial jamming. Nat Phys 14:613–620. doi:10.1038/s41567-0180109-9.

28. Park JA, Kim JH, Bi D, Mitchel JA, Qazvini NT, Tantisira K, Fredberg JJ. 2015. Unjamming and cell shape in the asthmatic airway epithelium. Nat Mater 14:1040–1048. doi:10.1038/nmat4357.

29. Sadati M, Qazvini NT, Krishnan R, Park CY, Fredberg JJ. 2013. Collective migration and cell jamming. Differentiation 86:121–125. doi:10.1016/j.diff.2013.02.003.

30. Mitchel JA, Das A, O’Sullivan MJ, Stancil IT, DeCamp SJ, Koehler S, Park JA. 2020. In primary airway epithelial cells, the unjamming transition is distinct from the epithelial-tomesenchymal transition. Nat Commun 11:5053. doi:10.1038/s41467-020-18841-7.

31. Foty RA, Steinberg MS. 2005. The differential adhesion hypothesis: a direct evaluation. Dev Biol 278:255–263. doi:10.1016/j.ydbio.2004.11.012.

32. Brodland GW. 2002. The differential interfacial tension hypothesis (DITH): a comprehensive theory for the self-rearrangement of embryonic cells and tissues. J Biomech Eng 124:188–197. doi:10.1115/1.1449491.

33. Canty L, Zarour E, Kashkooli L, François P, Fagotto F. 2017. Sorting at embryonic boundaries requires high heterotypic interfacial tension. Nat Commun 8:157. doi:10.1038/s41467-017-00146-x.

34. Rozman, J., Yeomans, J. M., & Sknepnek, R. (2023). Shape-tension coupling produces nematic order in an epithelium vertex model. Physical Review Letters, 131(22), 228301.

35. Muthukrishnan, S., Dewan, P., Tejaswi, T., Sebastian, M. B., Chhabra, T., Mondal, S., … & Vishwakarma, M. (2025). Glassy dynamics in active epithelia emerge from an interplay of mechanochemical feedback and crowding. bioRxiv, 2025s-11.

36. Martin, A. C. (2010). Pulsation and stabilization: contractile forces that underlie morphogenesis. Developmental biology, 341(1), 114–125.

37. Manning ML, Foty RA, Steinberg MS, Schoetz EM. 2010. Coaction of intercellular adhesion and cortical tension specifies tissue surface tension. Proc Natl Acad Sci U S A 107:12517–12522. doi:10.1073/pnas.1003743107.

38. Vogelstein, B., Papadopoulos, N., Velculescu, V. E., Zhou, S., Diaz Jr, L. A., & Kinzler, K. W. (2013). Cancer genome landscapes. science, 339(6127), 1546–1558.

39. Martincorena, I., & Campbell, P. J. (2015). Somatic mutation in cancer and normal cells. Science, 349(6255), 1483–1489.

40. Moruzzi, M., Nestor-Bergmann, A., Goddard, G. K., Tarannum, N., Brennan, K., & Woolner, S. (2021). Generation of anisotropic strain dysregulates wild-type cell division at the interface between host and oncogenic tissue. Current biology, 31(15), 3409–3418.

41. Sussman, D. M., Schwarz, J. M., Marchetti, M. C., & Manning, M. L. (2018). Soft yet sharp interfaces in a vertex model of confluent tissue. Physical review letters, 120(5), 058001.

42. Schindelin J, Arganda-Carreras I, Frise E, Kaynig V, Longair M, Pietzsch T, Cardona A. 2012. Fiji: an open-source platform for biological-image analysis. Nat Methods 9:676– 682. doi:10.1038/nmeth.2019.

43. Stringer C, Wang T, Michaelos M, Pachitariu M. 2021. Cellpose: a generalist algorithm for cellular segmentation. Nat Methods 18:100–106. doi:10.1038/s41592-020-01018-x.

44. Aigouy B, Umetsu D, Eaton S. 2016. Segmentation and quantitative analysis of epithelial tissues. Drosophila: Methods and Protocols 227–239. doi:10.1007/978-1-4939-63713_13.

45. Skamrahl M, Schünemann J, Mukenhirn M, Pang H, Gottwald J, Jipp M, Janshoff A. 2023. Cellular segregation in cocultures is driven by differential adhesion and contractility on distinct timescales. Proc Natl Acad Sci USA 120:e2213186120. doi:10.1073/pnas.2213186120.

46. Thielicke W, Sonntag R. 2021. Particle Image Velocimetry for MATLAB: Accuracy and enhanced algorithms in PIVlab. J Open Res Softw 9:12. doi:10.5334/jors.334.

47. Canty, L., Zarour, E., Kashkooli, L., François, P., & Fagotto, F. (2017). Sorting at embryonic boundaries requires high heterotypic interfacial tension. Nature communications, 8(1), 157.

48. Dye, N. A., Popović, M., Iyer, K. V., Fuhrmann, J. F., Piscitello-Gómez, R., Eaton, S., & Jülicher, F. (2021). Self-organized patterning of cell morphology via mechanosensitive feedback. Elife, 10, e57964.

